# Targeting interferon-_λ_ signaling promotes recovery from central nervous system autoimmunity

**DOI:** 10.1101/2021.08.17.456642

**Authors:** Sindhu Manivasagam, Jessica L. Williams, Lauren L. Vollmer, Bryan Bollman, Juliet M. Bartleson, Shenjian Ai, Gregory F. Wu, Robyn S. Klein

## Abstract

Type III interferons (IFNLs) are newly discovered cytokines, acting at epithelial and other barriers, that exert immunomodulatory functions in addition to their primary roles in antiviral defense. Here we define a role for IFNLs in maintaining autoreactive T cell effector function and limiting recovery in a murine model of multiple sclerosis (MS), experimental autoimmune encephalomyelitis (EAE). Genetic or antibody-based neutralization of the IFNL receptor (IFNLR) resulted in lack of disease maintenance during EAE, with loss of CNS Th1 effector responses and limited axonal injury. Phenotypic effects of IFNLR signaling were traced to increased antigen presenting cell (APC) function, with associated increase in T cell production of IFNγ and GM-CSF. Consistent with this, IFNL levels within lesions of CNS tissues derived from MS patients were elevated compared to MS normal appearing white matter (NAWM). Furthermore, expression of IFNLR was selectively elevated in MS active lesions compared to inactive lesions or NAWM. These findings suggest IFNL signaling as a potential therapeutic target to prevent chronic autoimmune neuroinflammation.

## Introduction

Multiple sclerosis (MS) is a debilitating T cell-mediated, demyelinating autoimmune disease of the central nervous system (CNS) (1) that progresses to severe disability in 90% of cases (2). Most MS patients initially present with relapsing-remitting MS (RRMS), in which episodes of neurological dysfunction are followed by partial or complete remission. Within twenty-five years, the majority of untreated RRMS patients progress to secondary progressive MS (SPMS) and experience increasing neurologic deterioration (3). The T helper (Th)1 cytokine, interferon gamma (IFN-γ), and the Th17 cytokine, interleukin (IL)-17, have demonstrated pathogenic roles in MS, while Th2 and regulatory T cells may limit disease activity (4, 5). Furthermore, T helper cells producing granulocyte-macrophage colony stimulating factor (GM-CSF) have been recently identified to be expanded in RRMS patient peripheral blood and cerebrospinal fluid (CSF) (6). The strong role of T cells in MS is further underscored by the number of T cell modulating therapies that have been successful for RRMS (7). However, currently approved immunomodulatory therapies are unable to prevent disease progression or promote recovery (8). Therefore, for patients with progressive disease, there are no treatments that prevent the chronic emergence of new disabilities or reverse deficits that previously developed. Studies indicate a critical role for CNS antigen presenting cells (APCs), including dendritic cells (DCs), in disease progression via the initiation and maintenance of T cell autoreactivity (9–12). These studies utilized the established murine model for MS, experimental autoimmune encephalomyelitis (EAE), to define roles for classical CD11c^+^MHC^+^ DCs in antigen presentation and epitope spreading to myelin-specific CD4^+^ T cells, which maintain disease. However, there are no therapies that specifically target these cell types or their cell-specific receptors.

Type III interferons (IFNs) or the lambda IFNs (IFNLs or IFN-λs), generate and sustain antiviral T cell responses via activation of its receptor, IFNLR, which is expressed by macrophages (13), subsets of DCs (14–17), and certain epithelial and endothelial barriers, including the blood-brain barrier (18). Type I and type III interferons are closely related, signaling through common JAK-STAT signaling pathways that lead to transcription of IFN-stimulated genes (ISGs) (19–21). Specifically, type I IFN binds to the IFNαβ receptor (IFNAR) and type III IFN binds to the heterodimeric receptor (IFNLR), which is comprised of two subunits, IFNLR1 and IL10Rβ (22). As IFNAR is ubiquitously expressed, type I IFN -inhibition of viral replication occurs in many cells types. In contrast, IFNLR-mediated antiviral responses exhibit specificity for viruses that replicate at barrier surfaces due to its cell-specific expression (21, 23). Studies in animal models of viral infections show that IFNL may act on DCs to modulate and augment downstream T cell polarization and effector function (15–17). Studies in tissues derived from patients with inflammatory disease or in murine models of these diseases indicate that IFNL contributes to inflammation and expansion of myeloid and T cell populations (24, 25), down-regulates Th2 cytokines and sustains Th1 activation (15, 26–28), including IFNγ production (29). However, the cellular targets of IFNL and mechanisms for regulation of CD4^+^ Th1 cell responses, especially in CNS autoimmune diseases, have not been defined.

Here, we demonstrate that IFNL signaling sustains myeloid cell-driven neuroinflammation in mice with experimental autoimmune encephalomyelitis (EAE), a model of the human disease MS. Induction of CNS autoimmunity in EAE relies on peripheral priming of CD4^+^ T cells, followed by local reactivation of effector myelin-specific T cells by CNS resident and infiltrating antigen presenting cells (APCs), including monocyte- and plasmacytoid-derived dendritic cells (30–32) and B cells (33). Using murine models of EAE, we show that IFNL promotes disease maintenance and axonal injury through maintenance of effector Th1 cells within the CNS. Using global and cell-specific deletion strategies, we demonstrate that, in the presence of IFNL, APCs maintain co-stimulatory molecules that participate in CD4^+^ T cell activation, with continued inflammatory cytokine production and disease maintenance. IFNL ligand and receptor levels are increased in MS patient lesions compared to MS normal appearing white matter (NAWM) and non-MS CNS tissues. Importantly, targeting IFNL with a single-dose neutralizing strategy is sufficient to reverse clinical effects and prevent axonal injury, identifying a potential new therapeutic target to treat MS.

## Results

### Loss of IFNL Signaling Reduces Severity of CNS Autoimmune Disease

To determine whether IFNL signaling impacts the induction of CNS autoimmunity, we induced EAE in WT and *Ifnlr1^−/−^* mice via active immunization with myelin oligodendrocyte glycoprotein (MOG)_35-55_ peptide. While day of EAE onset was similar (Fig. S1A), *Ifnlr1*^−/−^ animals developed significantly less severe EAE compared to WT animals at peak disease and subsequently maintained lower clinical activity scores during recovery (Fig. 1A, S1B). Consistent with the observed alterations in clinical disease course, areas containing ionized calcium binding adaptor molecule 1 (Iba1) expressing myeloid cells within lumbar spinal cord lesions were significantly decreased in *Ifnlr1*^−/−^ compared with WT mice at a chronic disease time-point (day 27 post immunization) (Fig. 1B). Importantly, *Ifnlr1*^−/−^ animals also exhibited decreased area of axonal injury compared to WT animals, as assessed by immunofluorescence (IF) detection of SMI-32 (non-phosphorylated neurofilament H) (Fig. 1B). No differences in levels of myelin basic protein (MBP), a protein found on the myelin sheath, or in damaged myelin basic protein (myelin basic protein_69-86_, dMBP) (Fig. S1C) were observed at this time-point. These results are consistent with published studies demonstrating that axonal damage may not require prior demyelination (34), and suggest a critical role for IFNL in promoting inflammation and axonal injury during CNS autoimmune disease.

**Figure 1.**
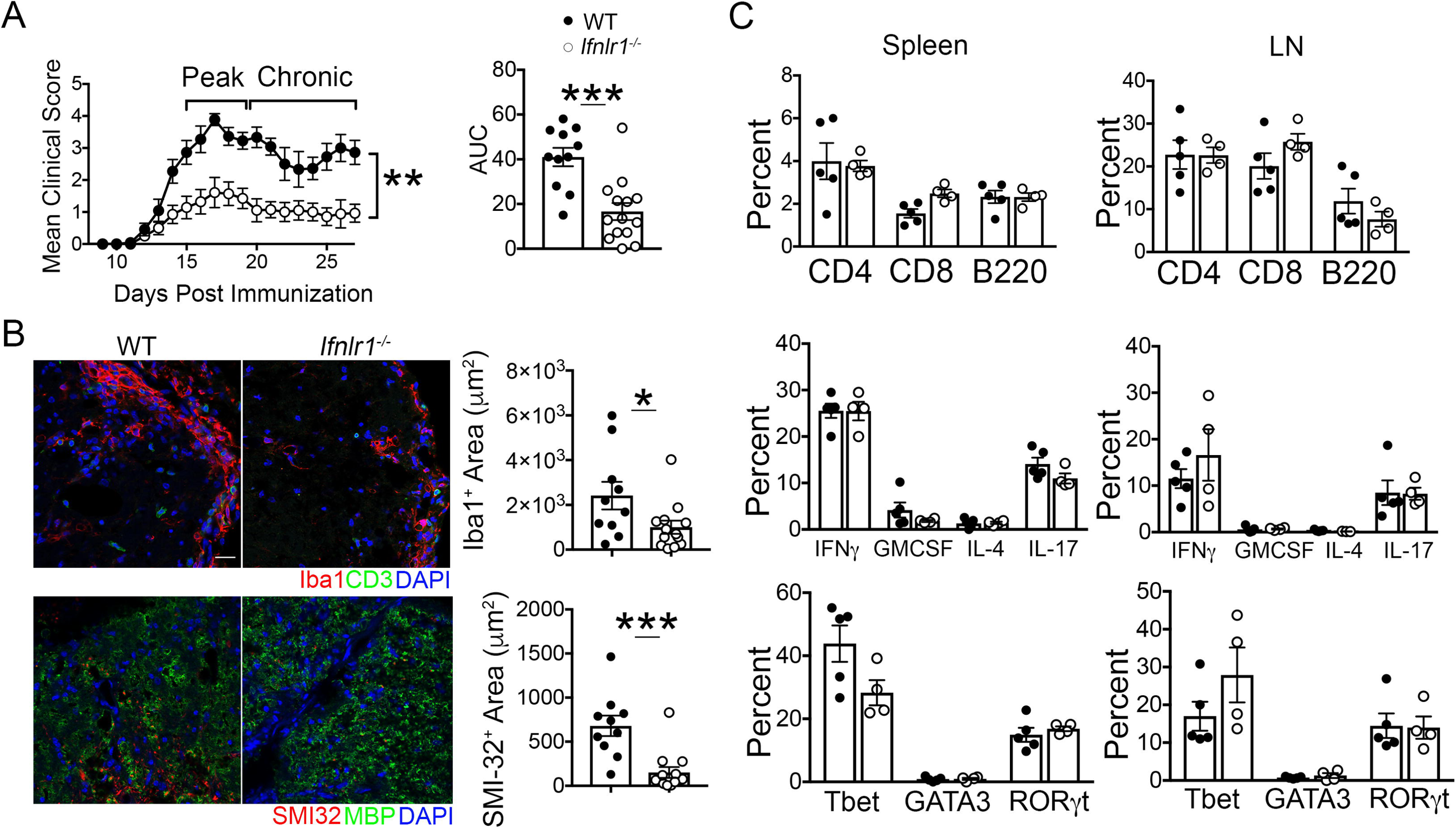
Loss of IFNL Signaling Reduces Severity of CNS Autoimmune Disease. **(A)** EAE clinical course and area under curve (AUC) analysis following active immunization with MOG_35-55_ peptide of WT and *Ifnlr1^−/−^* mice. Data are pooled from 2 independent experiments; n=11 WT and n=14 *Ifnlr1^−/−^* animals shown. **(B)** Immunofluorescence (IF) analysis of lesions within the ventral lumbar spinal cord of WT and *Ifnlr1^−/−^* mice during EAE at chronic timepoint indicated in (A). Iba1^+^ and SMI-32^+^ area were quantified. Data are pooled from 2 independent experiments; n=10 WT and n=14 *Ifnlr1^−/−^* animals shown. Scale bar = 20 μm. **(C)** WT and *Ifnlr1^−/−^* mice were actively immunized with MOG_35-55_ peptide. Cells were collected from spleen and draining lymph nodes 11 days following immunization. Flow cytometric analysis was performed on cells prior to stimulation to determine percentages of CD4^+^, CD8^+^, and B220^+^ cells. CD4^+^ cells were further gated on various cytokines and transcription factors; quantification is shown. Data are representative of 2 independent experiments; n=5 WT and n=4 *Ifnlr1^−/−^* animals shown. Data are presented as means ± SEM. *, *P* < 0.05; **, *P* < 0.01; ***, *P* < 0.001 by **(A)** Mann-Whitney *U* test and **(A-C)** two-tailed Student’s t test.

### IFNL Signaling Does Not Alter Peripheral Immune Responses after MOG_35-55_ Peptide Immunization

As baseline differences in peripheral immunity of WT and *Ifnlr1*^−/−^ animals could underlie differences in disease expression within the CNS, we examined percentages and phenotypes of lymphoid cells derived from the spleens and lymph nodes (LN) of naïve WT and *Ifnlr1^−/−^* mice after immunization with MOG_35-55_ peptide. Naïve animals showed similar percentages of splenic and LN CD4^+^, CD8^+^, and CD19^+^ cells in both genotypes (Fig. S2A-B). Analysis of peripheral lymphoid tissues 11 days following immunization with MOG_35-55_ also revealed no differences in percentages of CD4^+^, CD8^+^, and B220^+^ cells nor in percentages of CD4^+^ T cells expressing inflammatory cytokines or Th transcription factors (Fig 1C). Upon *in vitro* restimulation with MOG_35-55_, WT and *Ifnlr1*^−/−^ CD4^+^ Th1 polarized splenocytes and LN cells expressed similar levels of IFNγ and GMCSF (Fig. S2C), and induced EAE to similar extent following adoptive transfer to naïve, WT recipients (Fig. S2D). Together, these data suggest that loss of IFNL signaling does not significantly alter the initial peripheral immune response to MOG_35-55_ immunization, or the encephalitogenic capability of MOG_35-55_-specific T cells during EAE induction.

### IFNL Signaling is Required for the Maintenance of Autoimmune Neuroinflammation

Since peripheral T cell activation did not differ between WT and *Ifnlr1*^−/−^ animals, we examined whether IFNL signaling within the CNS differentially impacts T cell reactivation. Adoptive transfer of activated MOG_35-55_-specific, Th1 polarized WT CD4^+^ T cells into naïve WT and *Ifnlr1*^−/−^ recipients revealed no differences in time of EAE induction and peak clinical scores between the genotypes (Fig. S3A). However, recovery of neurologic function to baseline status occurred only in *Ifnlr1*^−/−^ recipients (Fig. 2A). Correspondingly, lesion areas and axonal injury, assessed as in Fig. 1, were decreased, while levels of detection of MOG and dMBP were unaffected in *Ifnlr1*^−/−^ compared to WT animals at a chronic disease time-point (day 22 post transfer) (Fig. 2B-D and S3B). Analysis of infiltrating and activated resident myeloid cells via detection of Iba1, which participates in membrane ruffling and phagocytosis (35), revealed decreased expression of the phagocytic marker CD68 in *Ifnlr1^−/−^* spinal cords compared to WT at the same time-point (Fig. 2C-D). To determine whether this decrease in Iba1^+^ cells was due to baseline differences, we examined naïve spinal cords and saw no differences in expression of Iba1 between WT and *Ifnlr1*^−/−^ animals (Fig. S4A). TUNEL assay analysis also demonstrated minimal co-localization with Iba1^+^ cells in all groups (Fig. S4B), suggesting differences in Iba1^+^ expression and activation are not due to cell death. Overall, these data support the notion that IFNL signaling maintains inflammation in CNS lesions during autoimmunity.

**Figure 2.**
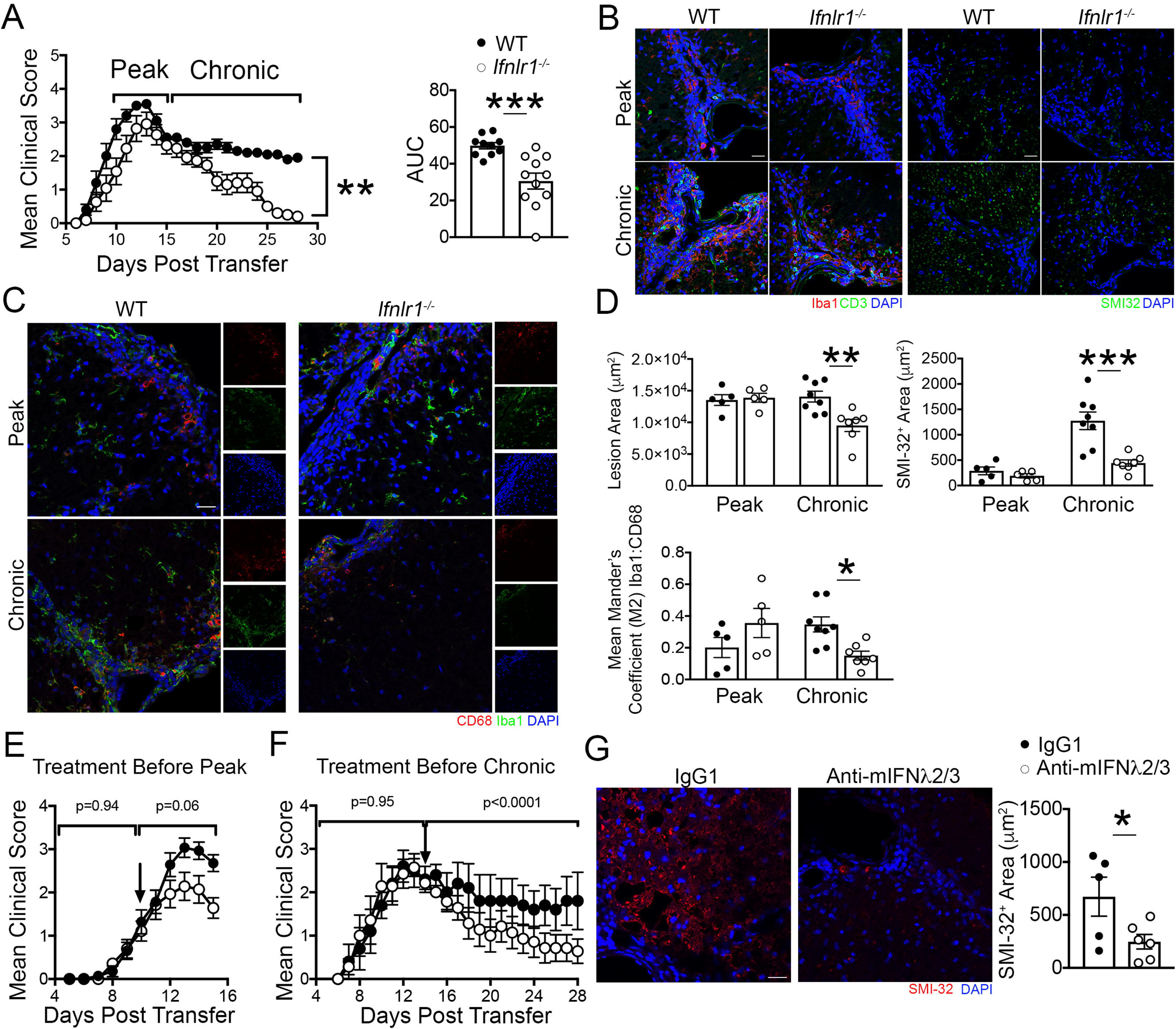
IFNL Signaling is Required for the Maintenance of Autoimmune Neuroinflammation. **(A)** Activated MOG-specific Th1 clones were injected into naïve WT or *Ifnlr1^−/−^* mice. Clinical course of recipient mice was monitored and AUC was quantified. Data are representative of 3 independent experiments; n=10 WT and n=11 *Ifnlr1^−/−^* animals shown. **(B-D)** IF analysis and quantification of lesions within the ventral lumbar spinal cord of WT and *Ifnlr1^−/−^* mice at the timepoints indicated in (A). Lesion area was delineated using CD3 and Iba1 staining and SMI-32^+^ area was quantified with comparisons between genotype. Mean Mander’s coefficient (M2) was used to quantify microglia and macrophage associated CD68 staining. Data are representative of 2 independent experiments; n=5 WT and n=5 Ifnlr1^−/−^ animals at peak timepoint and n=8 WT and n=7 Ifnlr1^−/−^ animals at chronic timepoint. Scale bar = 20 μm. **(E)** Activated, MOG_35-55_-specific WT Th1 clones were injected into naïve WT recipients. On day 10 post-transfer, neutralizing monoclonal mouse anti-mIFNL2/3 Ab (white circles) or an IgG1 control (black circles) was administered. Data are pooled from 3 independent experiments; n=14 animals per group shown. **(F)** Activated, MOG_35-55_-specific WT Th1 clones were injected into naïve WT recipients. On day 14 post-transfer, neutralizing monoclonal mouse anti-mIFNL2/3 Ab (white circles) or an IgG1 control (black circles) was administered. Data are representative of 3 independent experiments; n=5 IgG1 and n=7 anti-mIFNL2/3 Ab animals shown. **(G)** SMI-32^+^ area was quantified in ventral lumbar spinal cords of animals from (F). Data shown for n=5 IgG1 and n=6 anti-mIFNL2/3 Ab animals. Scale bar = 20 μm. Data are presented as means ± SEM. *, *P* < 0.05; **, *P* < 0.01; ***, *P* < 0.001 by **(A,E,F)** Mann-Whitney *U* test, **(A, G)** two-tailed Student’s t test, and **(D)** one-way ANOVA with multiple comparisons.

To evaluate the therapeutic potential of targeting IFNL, we determined whether antibody neutralization recapitulates results observed in *Ifnlr1*^−/−^ mice. Administration of monoclonal neutralizing antibody that targets murine IFNL2 and IFNL3 (anti-mIFNL2/3) versus non-specific IgG1 on day 10 following induction of EAE via adoptive transfer of MOG_35-55_ specific CD4^+^ T cells resulted in amelioration of peak EAE scores compared to controls (Fig. 2E). Similarly, single administration of anti-mIFNλ2/3 following peak EAE disease on day 14 post cell transfer, resulted in improved recovery from EAE (Fig. 2F) and decreased axonal injury (Fig. 2G). These studies not only validated data obtained through genetic approaches (Fig. 2A), but also suggest IFNL neutralization may be a therapeutic approach for reducing chronic neuroinflammation.

### IFNL Signaling Promotes Effector Function of Th1 Autoreactive Cells within the Inflamed CNS

The cytokines, interferon gamma (IFNγ), granulocyte-macrophage colony stimulating factor (GM-CSF), and interleukin (IL)-17, produced by effector CD4^+^ T cells within the inflamed spinal cord are strongly implicated in the induction and maintenance of EAE (36–38) and are elevated in postmortem human MS brain samples (39). Based on the crucial role of CD4^+^ T cell cytokine production in disease expression, we determined whether IFNLR signaling impacts the effector function of infiltrating spinal cord CD4^+^ T cells (Fig. S5A). Despite transfer of equal numbers of MOG_35-55_-specific WT Th1 cells, *Ifnlr1*^−/−^ recipients exhibited reduced numbers of CD4^+^ T cells within the spinal cord at peak disease compared to their WT counterparts (Fig. 3A-B). TUNEL assay analysis of spinal cords revealed increased CD3^+^ T cell death in *Ifnlr1*^−/−^ animals compared to WT animals (Fig. 3C), indicating IFNLR signaling may underlie T cell survival, and suggests a possible explanation for decreased T cell numbers. Examination of cytokine expression by spinal cord infiltrating CD4^+^ T cells at peak disease revealed that the majority expressed GM-CSF and IFNγ, but not IL-17 (Fig. 3D-E). There were significantly fewer GM-CSF^+^IFNγ^+^IL-17^-^ and GM-CSF^-^IFNγ^+^IL-17^-^ CD4^+^ T cells in the spinal cords of *Ifnlr1*^−/−^ compared to WT animals (Fig. 3E). Evaluation of CNS-derived CD4^+^ cells for expression of T cell transcription factors during peak EAE revealed decreased numbers of Tbet^+^RORγt^+^CD4^+^ cells in *Ifnlr1*^−/−^ versus WT animals. Analysis of other populations showed an increase in Tbet^-^RORγt^+^ CD4^+^ cells and a decrease in Tbet^+^RORγt^-^ CD4^+^ T cells in *Ifnlr1*^−/−^ versus WT animals (Fig. 3F). Analysis of T cell activation markers revealed decreased numbers of CD44^+^ and CD69^+^ CD4^+^ T cells derived from spinal cords of *Ifnlr1*^−/−^ mice with EAE compared with similarly affected WT animals, suggesting IFNLR signaling is required for complete Th1 cell effector function during EAE (Fig. 3G and S5B). Overall, these data indicate that loss of IFNL signaling significantly diminishes numbers of activated, autoreactive Th1 cells, consistent with reduced disease expression observed in *Ifnlr1^−/−^* mice.

**Figure 3.**
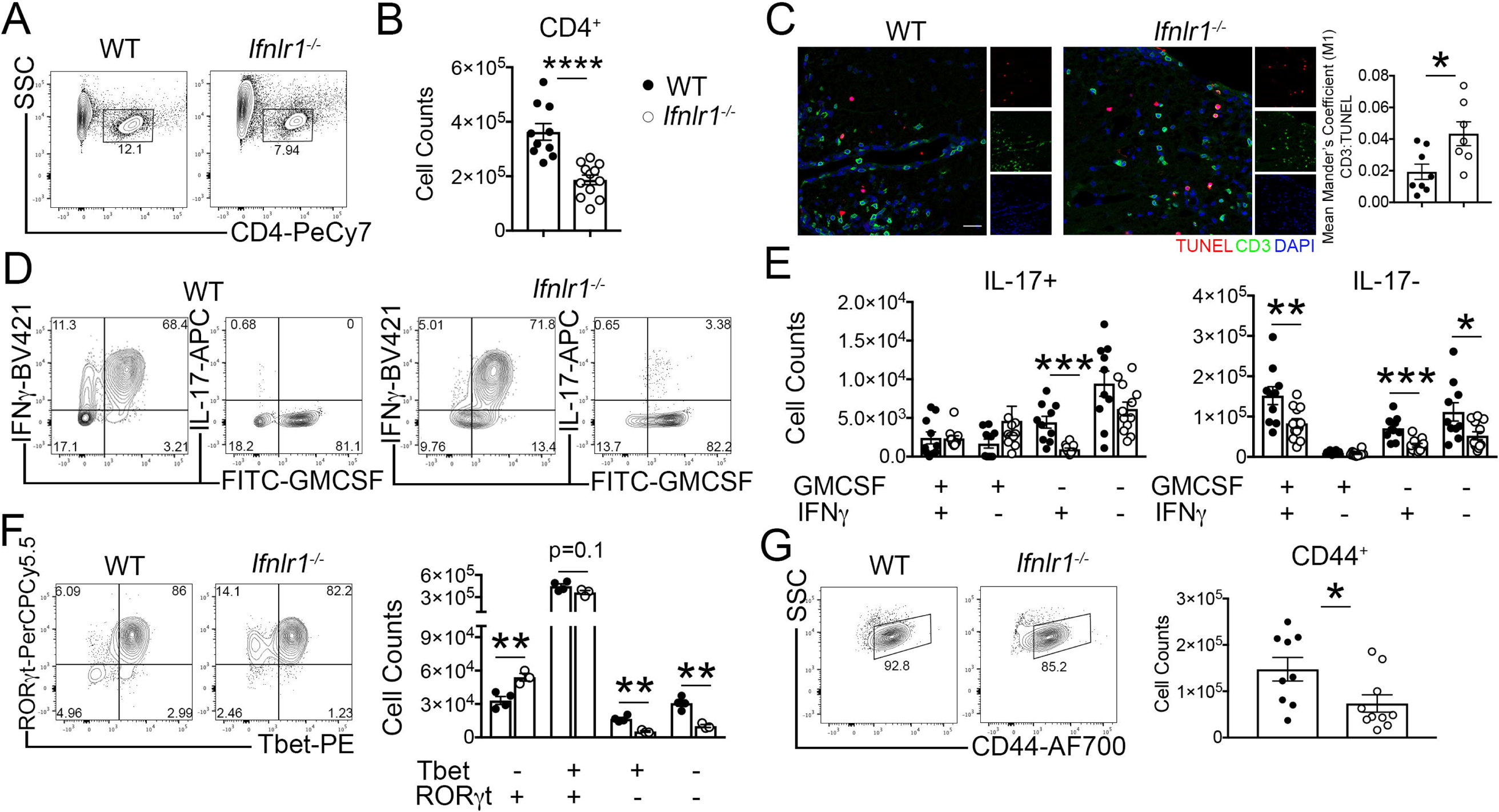
IFNL signaling maintains Th1 cell effector function during acute EAE. **(A-B)** Activated MOG-specific Th1 clones were injected into naïve WT or *Ifnlr1^−/−^* mice. Flow cytometric analysis of infiltrating cells was performed on lumbar spinal cords during peak EAE (Fig. 2A). CD4^+^ cells were identified from a live, single cell gate and quantified. Data are pooled from 2 independent experiments; n=10 WT and n=12 *Ifnlr1^−/−^* animals shown. (**C**) Immunofluorescence analysis of TUNEL^+^ and CD3^+^ co-localization in lumbar spinal cords at chronic EAE timepoint (Fig. 2A). Data shown for n=8 WT and n=7 *Ifnlr1^−/−^* animals. Scale bar = 20 μm. **(D-E)** Of the CD4^+^ cells shown in (B), GMCSF^+^, IFNγ^+^ and IL-17^+^ cells were identified and quantified. Data are pooled from 2 independent experiments; n=10 WT and n=12 *Ifnlr1^−/−^* animals shown. **(F)** CD4^+^ cells were gated for Tbet and RORγt and quantified. Data are representative of 2 independent experiments; n=4 WT and n=3 *Ifnlr1^−/−^* animals shown. **(G)** CD4^+^ cells were gated on CD44 and quantified. Data are pooled from 2 independent experiments; n=9 WT and n=10 *Ifnlr1^−/−^* animals shown. Data are presented as means ± SEM. *, *P* < 0.05; **, *P* < 0.01; ***, *P* < 0.001; ****, *P* < 0.0001 by two-tailed Student’s *t* test, comparing WT vs *Ifnlr1^−/−^*.

### IFNL Signaling Promotes Myeloid Cell Activation within the CNS

As intrinsic IFNLR signaling did not affect T cell encephalitogenicity, we next examined the impact of IFNLR deficiency on the phenotypes and functions of CNS myeloid subsets during EAE (Fig. S5A). We found that numbers of CD11b^+^ and CD11c^+^ cells derived from spinal cords of *Ifnlr1*^−/−^ animals compared to WT controls were significantly decreased (Fig. 4A). Within these populations, numbers of cells expressing costimulatory molecule, CD86, which is required for effective local CD4^+^ T cell reactivation during EAE, were also significantly decreased in *Ifnlr1*^−/−^ versus WT mice (Fig. 4B). Furthermore, analysis of median fluorescence intensity (MFI) of CD86 per cell revealed diminished expression in *Ifnlr1^−/−^* compared to WT in CD11b^+^CD11c^+^ and CD11b^+^CD11c^-^ populations (Fig. 4C). These data suggest that in the absence of IFNLR1, there are fewer activated APCs to provide the costimulatory signals necessary for proper T cell reactivation within the inflamed CNS.

**Figure 4.**
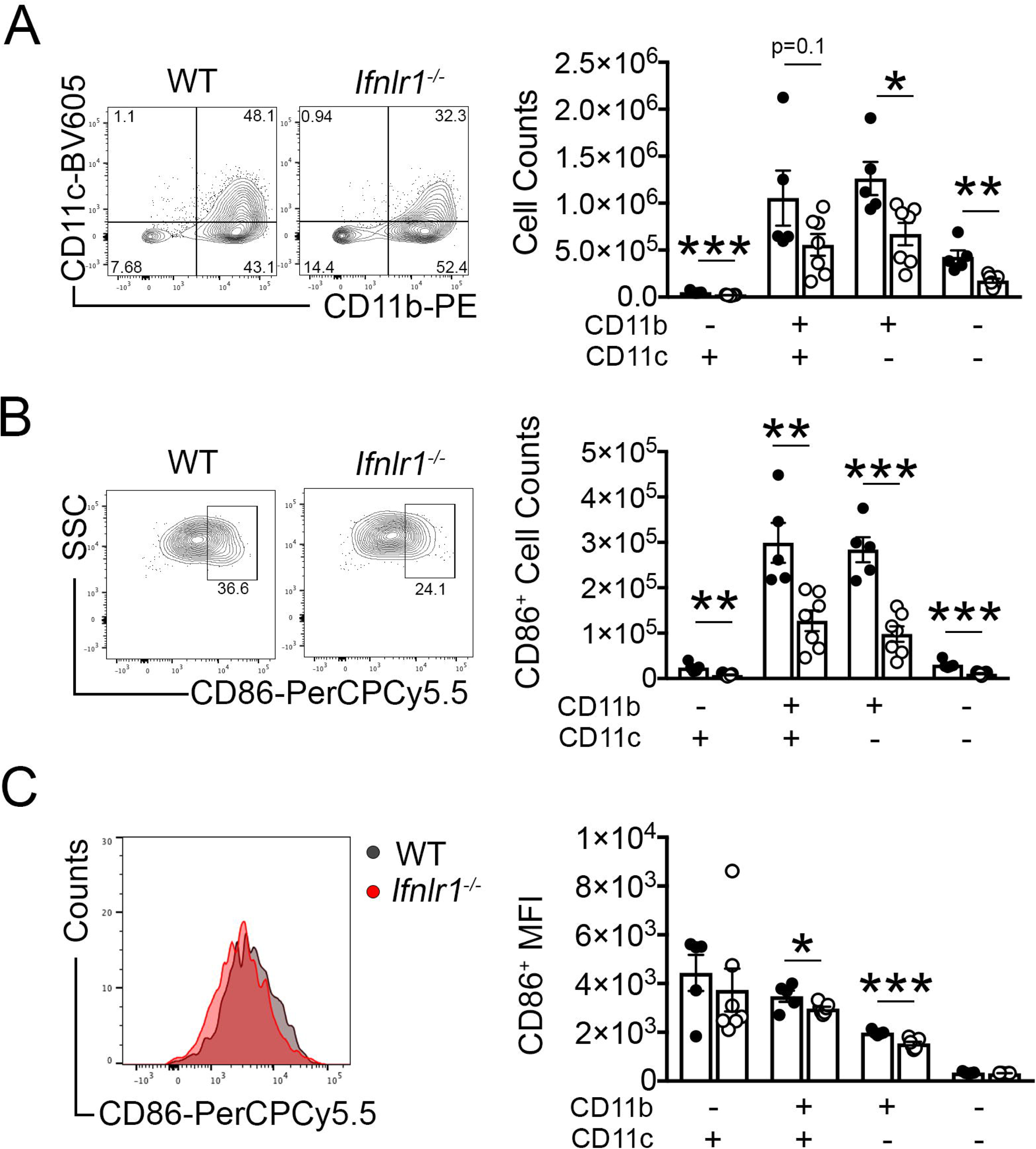
IFNL signaling maintains myeloid cells in the CNS during acute EAE. **(A)** Live, single cells were gated on CD45, followed by CD11b and CD11c gating. Numbers of these populations were quantified. Each of these populations was analyzed for **(B)** numbers of cells expressing CD86 and **(C)** median fluorescence intensity (MFI) of CD86. Representative plots shown for the CD11b^+^CD11c^+^ population in (B-C). Data are shown for n=5 WT and n=7 *Ifnlr1^−/−^* animals. Data are presented as means ± SEM. *, *P* < 0.05; **, *P* < 0.01, ***, *P* < 0.001 by two-tailed Student’s *t* test, comparing WT vs *Ifnlr1^−/−^*

### IFNL Signals Through CD11c^+^ Cells to Maintain Neuroinflammation

Consistent with prior studies detecting IFNLR expression by DCs (15, 17), *Ifnlr1* mRNA within spinal cords at peak EAE co-localized with CD11c mRNA within cells in perivascular lesions, as well as in the parenchyma (Fig. 5A). As shown previously (18), *Ifnlr1* mRNA expression was also detected in microvascular endothelial cells (Fig. 5A). Based on observed alterations in myeloid cell numbers in WT and *Ifnlr1*^−/−^ animals (Fig. 4), and prior studies demonstrating both macrophages (13) and dendritic cells (17) are IFNL targets, we determined the cell specific effects of IFNL signaling in EAE via MOG_35-55_ peptide immunization of *Ifnlr1*^fl/fl^CD11c-Cre (targeting dendritic cells) (40) and *Ifnlr1*^fl/fl^LysM-Cre (targeting neutrophils and macrophages) (41) animals. *Ifnlr1*^fl/fl^CD11c-Cre^+^ animals exhibited significant improvement of EAE scores in the chronic disease phase compared to *Ifnlr1*^fl/fl^CD11c-Cre^-^ animals (Fig. 5B), while no differences were observed between *Ifnlr1*^fl/fl^LysM-Cre^+^ and *Ifnlr1*^fl/fl^LysM-Cre^-^ animals (Fig. S6C). *Ifnlr1*^fl/fl^CD11c-Cre^+^ animals also exhibited significant decreases in axonal injury, but limited differences in Iba1^+^ or MBP^+^ cells compared with *Ifnlr1^fl/fl^*CD11c-Cre^-^ animals (Fig. 5C, S6D). These data suggest that IFNLR1 expression by CD11c^+^ cells influences EAE disease course and neuropathology.

**Figure 5.**
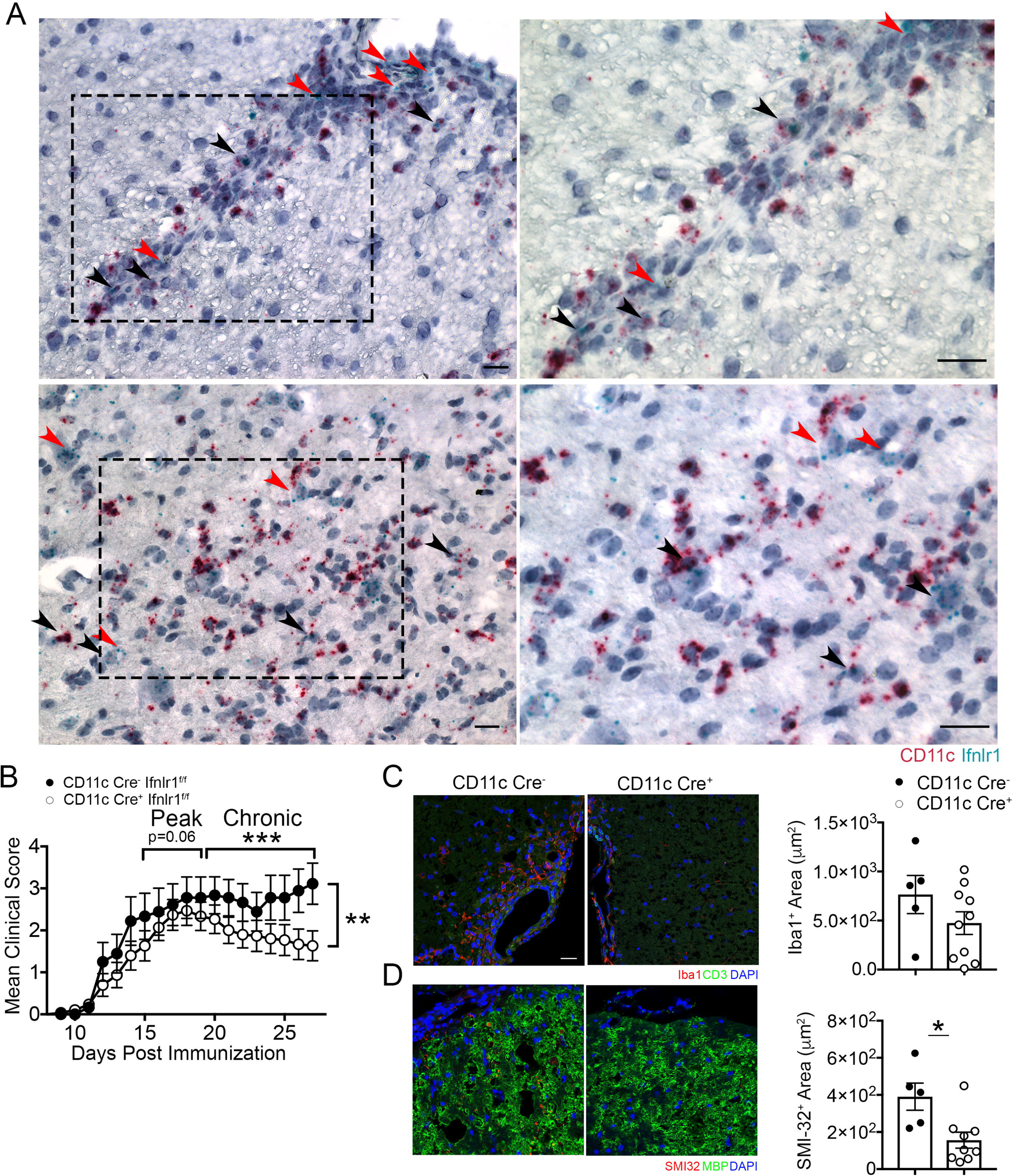
CD11c^+^ cells are critical targets of IFNL during EAE. **(A)** Representative images of dual *in situ* hybridization for *Ifnlr1* and CD11c RNA on WT spinal cords collected at peak EAE. Black arrowheads demonstrate co-localization of *Ifnlr1* and *CD11c*. Red arrowheads depict *Ifnlr1^+^CD11c^-^* cells. **(B)** *Ifnlr1^fl/fl^*CD11c-Cre^-^ and *Ifnlr1^fl/fl^*CD11c-Cre^+^ mice were monitored for EAE disease course following active immunization. Data are pooled from 3 independent experiment; n=9 *Ifnlr1^fl/fl^*CD11c-Cre^-^ and n=15 *Ifnlr1^fl/fl^*CD11c-Cre^+^ animals shown. IF analysis of **(C)** Iba1^+^ and **(D)** SMI-32^+^ area in ventral lumbar spinal cord of *Ifnlr1^fl/fl^*CD11c-Cre^-^ and *Ifnlr1^fl/fl^*CD11c-Cre^+^ mice at 27 days post immunization. Scale bar = 20 μm. Data shown for (C) n=5 *Ifnlr1^fl/fl^*CD11c-Cre^-^ and n=10 *Ifnlr1^fl/fl^*CD11c-Cre^+^ animals and (D) n=5 *Ifnlr1^fl/fl^*CD11c-Cre^-^ and n=9 *Ifnlr1^fl/fl^*CD11c-Cre^+^ animals. Data are presented as means ± SEM. *, *P* < 0.05; **, *P* < 0.01, ***, *P* < 0.001 by **(B)** Mann-Whitney *U* test and by **(C-D)** two-tailed Student’s *t* test.

### IFNL Ligand and Receptor Levels are Elevated in MS Lesions

Given that IFNL signaling is critical for the maintenance of encephalomyelitis in mice, we wondered if IFNL expression differed within inflammatory demyelinating lesions versus normal appearing white matter (NAWM) within CNS tissues of MS patients. Immunohistochemical detection of IFNL in human post-mortem specimens derived from the brains of patients with no neurologic diseases (non-MS), RRMS, and SPMS patients (Table 1 and Fig. 6A) revealed similar levels in the NAWM of MS patients and non-MS controls. However, CNS lesions in tissues derived from SPMS patients exhibited increased expression of IFNL compared to NAWM of SPMS patients. There were no significant differences in level of IFNL expression between lesions of RRMS and SPMS patients (Fig. 6B). In RRMS patients, IFNL was localized to vessels while in SPMS patients IFNL was also observed in parenchymal cells (Fig. 6A). Quantitative RT-PCR detection of *Ifnlr1* mRNA, within human post-mortem CNS specimens derived from non-MS and MS patients (Table 2), revealed significantly increased levels of *Ifnlr1* mRNA in active MS lesions, compared to non-MS tissue, MS NAWM, and MS inactive lesions (Fig. 6C). These data, in conjunction with our murine findings, suggest IFNL signaling plays a critical role in promoting and maintaining inflammation within demyelinating lesions during CNS autoimmune disease.

**Figure 6.**
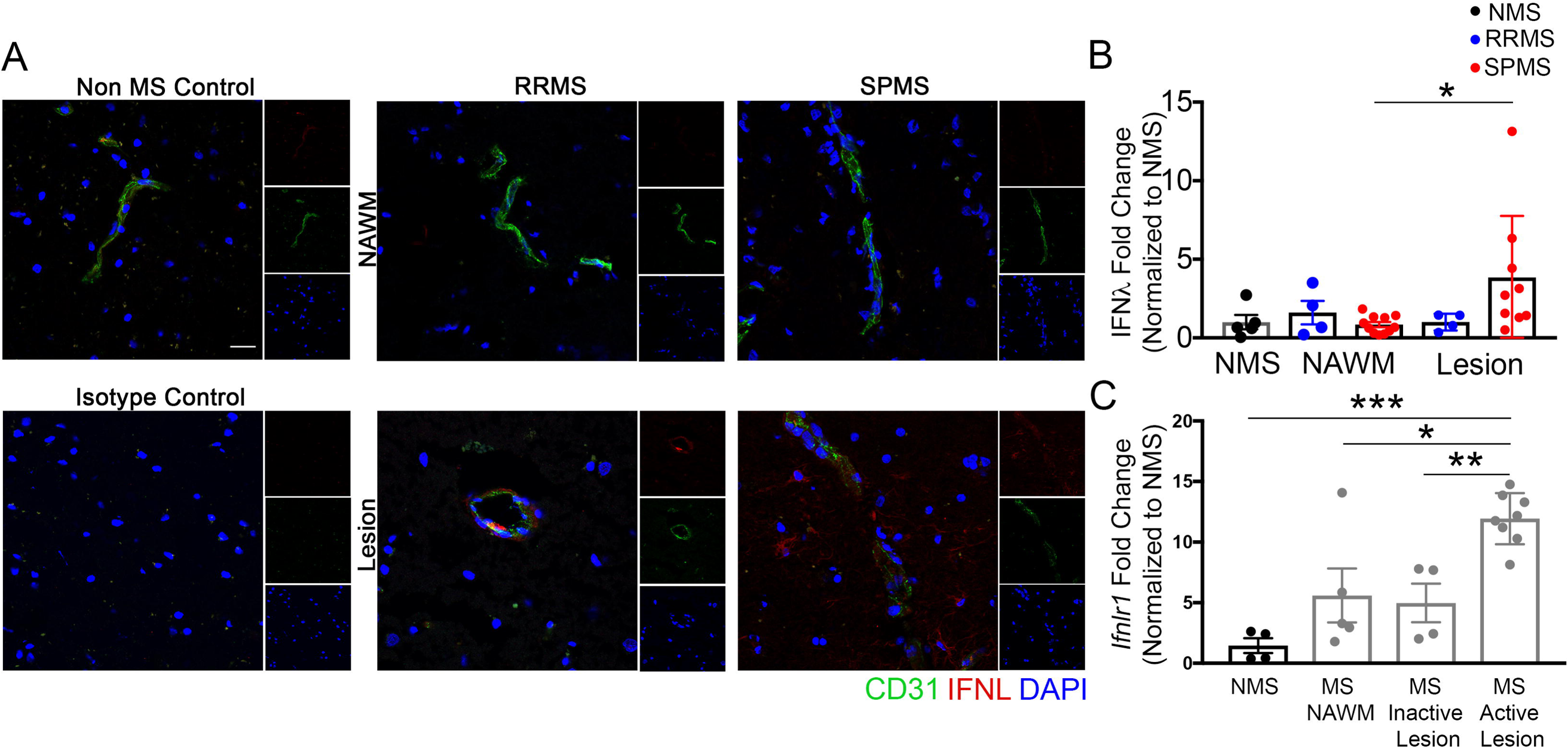
IFNL and IFNLR1 expression are increased in MS lesions. **(A)** Postmortem tissue specimens were analyzed: n= 5 samples from 5 non-MS (NMS) controls, n = 4 lesion samples and n=4 NAWM samples from 3 RRMS patients, and n=9 lesion samples and n=11 NAWM samples from 7 SPMS patients (Table 1). Non-MS patient samples were obtained from cortical white matter while MS patient samples were obtained from multiple CNS regions including the cortex, putamen, periventricular white matter and midbrain. Perivascular and parenchymal localization of IFNL in normal appearing white matter (NAWM) and in lesions were visualized using IF for CD31 and IFNL. Nuclei were counterstained with DAPI. Scale bar = 20 μm. **(B)** Area of IFNL was quantified. All values were normalized to the average of the NMS samples; each data point represents one tissue sample. **(C)** RNA was analyzed from postmortem human specimens: n=4 samples from 3 NMS controls and n= 5 NAWM samples, n= 4 inactive lesion samples, n= 8 active lesion samples from 18 MS patients (Table 2). Relative *Ifnlr1* mRNA expression was measured by qRT-PCR. All values were normalized to the average of the NMS samples; each data point represents one tissue sample. Data are presented as means ± SEM. *, *P* < 0.05; **, *P* < 0.01, ***, *P* < 0.001 by **(B-C)** one-way ANOVA with multiple comparisons.

**Table 1.**
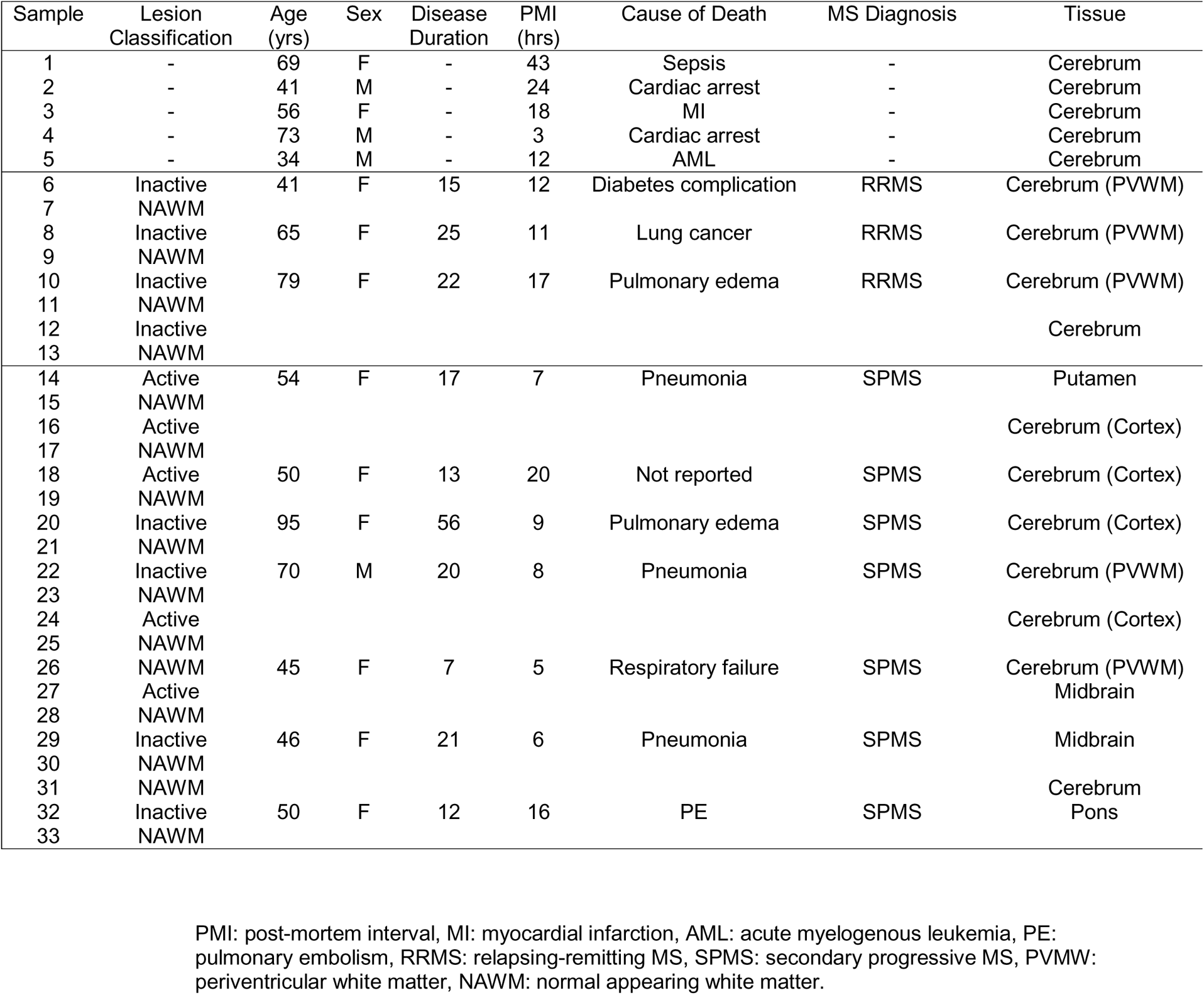
Demographics of post-mortem human samples for immunofluorescence analysis.

**Table 2.**
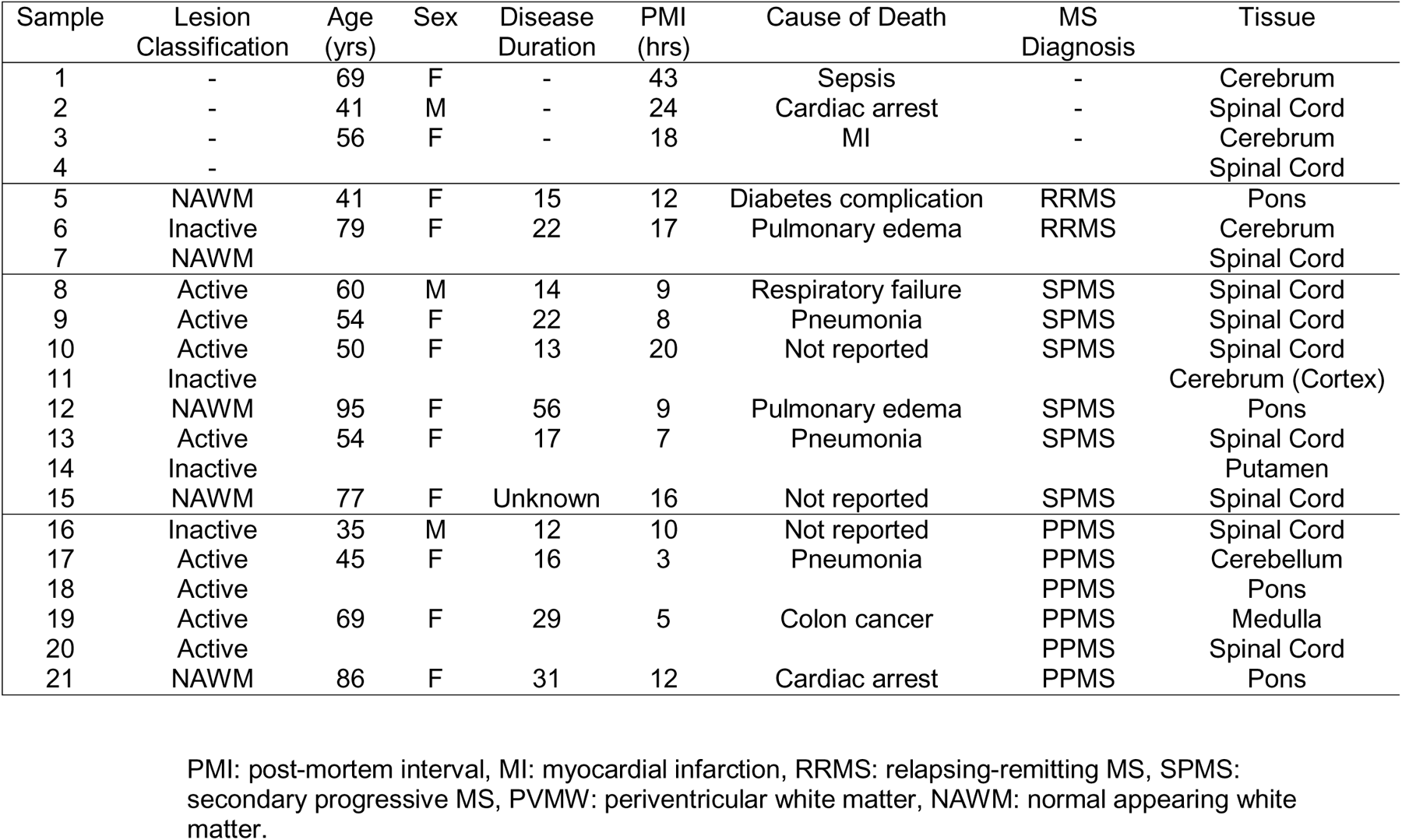
Demographics of post-mortem human samples for qRT-PCR analysis.

## Discussion

The identification of new therapeutic targets is essential for the development of new drugs to prevent chronic disease in MS patients. Our study identifies IFNL-IFNLR expression during CNS autoimmunity as a critical component in the maintenance and function of autoreactive T cells. Inactivation of IFNL signaling, via targeted deletion of *Ifnlr1*, led to reduced axonal pathology and recovery of neurologic function. These phenotypes correlated with reduced numbers of CNS infiltrating T cells and myeloid cells. The effects of IFNLR1 signaling were T cell extrinsic, as *in vitro* restimulated Th cells from WT and *Ifnlr1^−/−^* animals induced similar EAE following adoptive transfer into WT recipients. Instead, IFNL targeted myeloid cells, as CD11c^+^ cell-specific deletion of IFNLR1 improved recovery following EAE. IFNL ligand and receptor levels are also increased in MS patient lesions compared to NAWM. Importantly, neutralization of IFNL with a single administration of anti-mIFNλ disease and improved recovery when given before or after peak EAE, respectively. Overall, our study highlights IFNL as a potential therapeutic target to prevent chronic neuroinflammation.

In concordance with our findings, recent studies highlight that IFNL plays a significant role in immune-driven diseases. In a murine model of TLR7-induced lupus, IFNL promotes immune dysregulation through expansion of myeloid and T cell populations in the spleen and blood and induction of chemokine production by keratinocytes (24). Furthermore, IFNL protein levels are upregulated in patients with rheumatoid arthritis (42–45) and contribute to inflammation and cartilage degradation during osteoarthritis (25). Studies of allergic airway disease, vaccination induced inflammation, and *in vitr*o models examining T cell subtypes reveal the ability of IFNL to inhibit Th2 polarization (26–28, 46) and promote IFNγ production (15, 27, 28, 47), indicating that IFNL may enhance Th1 function (48). The mechanisms by which IFNL alters T cell function are yet to be fully elucidated but our data indicate that T cells are not directly targeted. Additionally, in corroboration with our work, studies of viral influenza suggest that IFNL modulation of CD11c^+^ DC responses affects the downstream adaptive immune response (17, 49). Conventional DCs may also produce IFNL to promote anti-tumor immunity via promotion of a Th1 microenvironment (50). One study has also indicated that IFNL may inhibit IL-17 responses (51), which may explain the increase in RORγt cells observed in *Ifnlr1*^−/−^ animals compared to WT animals in the current study (Fig. 3F). IFNL may also alter macrophage (13), natural killer cell (52), and neutrophil (51, 53, 54) polarization and cytokine production, as well as B cell differentiation (55).

Examination of IFNL within the CNS has primarily been in the context of neuroinvasive viral illnesses. During West Nile virus (WNV) encephalitis, IFNL substantially limits neuroinvasion by stabilizing the blood brain barrier (18, 56). In our present work, lack of IFNL signaling reduces numbers of infiltrating T cells within the CNS and ameliorates EAE. These differences are likely because T cells and viruses utilize different mechanisms to enter the CNS. While WNV peripheral infection relies on blood-brain barrier disruption to cross into the CNS (57, 58), activated leukocytes are able to bind to adhesion molecules upregulated on the endothelium and extravasate in a transcellular fashion across the barrier (59). Aside from the endothelium, IFNL also acts on other CNS cell types; IFNL treatment of *in vitro* culture of primary astrocytes and neurons results in diminished replication of herpes simplex virus type I within these cells (60). These studies focused on the effect of IFNL during acute viral infection, particularly on induction of JAK/STAT mediated antiviral gene transcription and control of replication. Our findings suggest IFNL may have additional immunomodulatory roles within the CNS, especially in the context of ongoing inflammation. The role of IFNL in chronic neuroinvasive viral infection is unknown but may be similar to our observations, especially since IFNL has been shown to exert opposing effects on T cells based on viral chronicity in a model of lymphocytic choriomeningitis virus (61).

Non-redundant roles that distinguish IFNL functions from those of type I IFNs have undergone limited investigation. Models of epithelial viral infections of lung and gut have shown temporal and spatial differences between type I and type III IFN responses (62, 63). One explanation for these differences is that while IFNAR is expressed ubiquitously, IFNLR is only expressed on a limited set of cells (64, 65). Our data also indicate that type I and type III IFN responses are quite distinct during CNS autoimmunity. While we show that *Ifnlr1*^−/−^ mice recover from EAE, *Ifnar1*^−/−^ mice develop a more severe disease course, a phenotype mediated by IFNAR signaling in myeloid cells (66). Furthermore, type I IFN may skew the immune response away from Th1 and towards Th2 (67–69), while multiple studies, including ours, show that type III promotes Th1 polarization. This dichotomy demonstrates the protective function of type I IFN and inflammation-promoting properties of type III IFN during CNS autoimmunity.

Overall, our results highlight the important role that myeloid cells play in Th1 cell mediated maintenance of neuroinflammation that requires IFNL signaling (70, 71). Furthermore, our antibody mediated neutralization studies highlight the therapeutic potential of targeting IFNL since neutralization, even after establishment of encephalomyelitis, was significantly beneficial. Further studies identifying methods to reduce prolonged inflammation and damage in the CNS are necessary for developing therapeutics that prevent the chronic disease associated with MS.

## Methods

### Animals

C57BL/6 (The Jackson Laboratory) and *Ifnlr1*^−/−^ mice (Bristol-Myers Squibb) (18, 72) were maintained in specific pathogen-free conditions at Washington University in St. Louis. Littermate *Ifnlr1^+/+^*, *Ifnlr1^+/−^*, *Ifnlr1^−/−^* co-housed animals were bred and maintained in specific pathogen-free conditions at Washington University in St. Louis. *Ifnlr1^fl/fl^* CD11c-Cre and *Ifnlr1^fl/fl^* LysM-Cre lines were obtained from Dr. Megan Baldridge at Washington University in St. Louis) (73). All animal studies were performed in accordance with the Animal Care and Use Committee guidelines of the National Institutes of Health and were conducted under protocols approved by the Animal Care and Use Committee of Washington University School of Medicine.

### EAE induction

Active EAE was induced in 8-10 week old male and female mice by subcutaneous immunization with murine MOG_35-55_ peptide (Genscript) and complete Freund’s adjuvant, as previously described (74). Pertussis toxin (List Biological Labs) (400ng/mouse) was administered intraperitoneally on days 0 and 2 following immunization. Animals were monitored blindly for weight loss and clinical scores based on the following EAE scale: 1, tail weakness; 2, difficulty righting; 3, one hindlimb paralysis; 4, two hindlimb paralysis; 5, moribund or dead. For adoptive transfer experiments, spleens from immunized animals were used to generated WT or *Ifnlr1*^−/−^ MOG_35-55_-specific Th1 clones, as previously described (75) and 10^7^ cells were transferred to naïve recipients via retroorbital injection. The group size for EAE studies was determined on the basis of previous study reports and our past experience using this model.

### Murine CNS tissue immunofluorescence (IF)

Frozen sections preparation and detection of cell markers were accomplished as previously described (76). Briefly, mice were anesthetized and perfused with ice-cold Dulbecco’s Phosphate Buffered Saline (dPBS), followed by 4% paraformaldehyde (PFA). Spinal cords were fixed overnight in 4% PFA, and then cryoprotected in 30% sucrose at 4°C for 72 hours; sucrose was exchanged every 24 hours during that time. Tissues were frozen in OCT (Fisher) and sectioned into 10μm sections. Tissue sections were blocked with goat or donkey serum and 0.1% Triton X-100 (Sigma-Aldrich) for 1 hour at room temperature. They were next exposed to anti-CD3 (1:200, BD, Catalog 555273), -Iba1 (1:250, Wako, Catalog 019-19741), -MOG (1:80, R&D Systems, AF2439), -MBP (1:100, Abcam, ab40390), -dMBP (1:2000, Millipore, Catalog AB5864), -SMI-32 (1:1000, BioLegend, Catalog 801702), or -CD68 (1:500, Bio-Rad, Catalog MCA1957) overnight at 4°C. Secondary antibodies conjugated to Alexa 488, Alexa 555, or Alexa 647 (1:400, Invitrogen) were applied for 1 hour at room temperature. Nuclei were counterstained with 4,6-diamidino-2-phenylindole (DAPI; Invitrogen). Coverslips were applied with ProLong Gold Anti Fade Mountant (Thermo Fisher).

Terminal deoxynucleotidyl transferase dUTP nick end labeling (TUNEL) was performed on sections using the In Situ Cell Death Detection Kit, TMR Red (Millipore Sigma), according to manufacturer’s protocol. Immunofluorescent images were acquired using the 20x or 40x objective of a Zeiss LSM 880 confocal laser scanning microscope. The mean positive area was determined using appropriate isotype control antibodies and quantified using Volocity (Perkin Elmer) and Fiji (NIH) (77) image analysis software. Co- localization analysis was performed using Just Another Colocalization Plugin (JACoP) in Fiji.

### *In situ* hybridization

Frozen sections were prepared as described under the murine CNS tissue IF section. Following sectioning of tissue, RNAscope 2.5 HD Duplex Assay (Advanced Cell Diagnostics) and RNAscope 2.5 HD Assay – Red (Advanced Cell Diagnostics) were performed as per manufacturer’s instructions. Probes (Advanced Cell Diagnostics) against *Ifnlr1* and CD11c *(Itgax)* were used. Following *in situ* hybridization, tissues were counterstained with 50% hematoxylin in water. Images were acquired using the 20x, 40x, or 63x objective of a Zeiss Cell Observer Inverted Microscope.

### CNS leukocyte isolation and flow cytometric analysis

Following cardiac perfusion with PBS, cells were isolated from the spleen, lymph node (LN) and spinal cords of WT or *Ifnlr1*^−/−^ mice. Spleen and LN were pushed through 70 μ M strainer to create a single cell suspension, following which cells from the spleen were incubated with ACK lysing buffer (Thermo Fisher Scientific). CNS tissue was digested in HBSS (Gibco) containing 0.05% collagenase D (Sigma), 0.1 μg/mL TLCK trypsin inhibitor (Sigma), 10 μg/mL DnaseI (Sigma) and 10mM Hepes pH 7.4 (Gibco) for 1 h at 37 degrees C. Tissue was pushed through 70 μM strainer to create a single cell suspension, and then centrifuged at 500 x *g* for 10 min. Cell pellets were resuspended in 37% Percoll (GE Healthcare) and centrifuged at 1200 x *g* for 30 min with deceleration set to 0, in order to remove myelin debris. Cell pellet was resuspended in PBS, counted using hemocytometer, and stained at a concentration of 1:200 with fluorescently conjugated antibodies. The following antibodies were used: CD4 (Clone RM4-5, BioLegend, Cat 100527), GMCSF (Clone MP1-22E9, BioLegend, Cat 505403), IFNγ (Clone XMG1.2, BioLegend, Cat 505829), IL-17 (Clone TC11-18H10.1, BioLegend, Cat 506915), Tbet (Clone 4B10, BD Biosciences, Cat 561265), RORγt (Clone Q31-378, BD Biosciences, Cat 562683), CD44 (Clone IM7, BioLegend, Cat 103025), CD69 (Clone H1.2F3, BioLegend, Cat 104512), CD11b (Clone M1/70, BD Biosciences, Cat 557397), CD45 (Clone 30-F11, BioLegend, Cat 103113), CD11c (Clone N418, BioLegend, Cat 117333), CD86 (Clone GL-1, BioLegend, Cat 105026), and CD8 (Clone 53-6.7, BioLegend, Cat 100729) as previously described (78). Live/Dead Fixable Aqua Dead Cell Stain Kit (Thermo Scientific) was used to identify live cell populations. Data were collected using a Fortessa X20 flow cytometer (BD) or LSR-II flow cytometer (BD) and analyzed using FlowJo software. Percentage data from flow cytometer was multiplied with hemocytometer cell counts to obtain a total number of cells per tissue.

### In vivo administration of anti-IFNL2/3 neutralizing antibody

Mice used in the neutralization studies were randomly assigned to a treatment group on day 10 or day 14 post-Th1 cell transfer, anesthetized with 2% isoflurane, and administered either 100 μg of neutralizing monoclonal mouse anti-mIFNL2/3 or 100 μg of a monoclonal mouse antibody specific for *E. coli* β-galactosidase (InvivoGen) in 250 μl sterile PBS via retroorbital injection. Clinical scores and weight loss were monitored blindly.

### Postmortem human tissue immunofluorescence analysis

Postmortem CNS tissue from non-MS controls and patients with clinically defined MS were obtained from The Neuroinflammatory Disease Tissue Repository at Washington University in St. Louis. Frozen sections were hydrated and blocked in 0.1% Triton X-100 and 10% donkey serum, followed by incubation with mouse anti-human CD31 (1:20, BD, Catalog 550389), and goat anti-human IFNL (1:25, R&D Systems, Catalog AF1587) antibodies. Secondary antibodies conjugated to Alexa 488 or Alexa 555 (1:400, Molecular Probes) were applied for 1 h at room temperature. Nuclei were counterstained with DAPI (Molecular Probes). Immunofluorescent images were acquired in a blinded manner using the 20x or 40x objective of a Zeiss LSM 880 confocal laser scanning microscope. The mean positive area was determined using appropriate isotype control antibodies and quantified using Fiji (NIH) (77) image analysis software

### Quantitative reverse transcription PCR (qRT-PCR)

RNA was extracted using the RNeasy Micro Kit (Qiagen). RNA was treated with DNase and cDNA was synthesized using Multiscribe reverse transcriptase (Applied Biosystems). qRT-PCR was performed using *Power* SYBR® Green PCR master mix (ThermoFisher), primers specific for human *Ifnlr1* (79) and *Gapdh,* and CFX 384 Touch Real-Time PCR Detection System (Biorad). CT values greater than 40 were not detected and therefore, not quantified. The delta delta CT method was applied to determine differences in gene expression levels after normalization to *Gapdh*.

### Statistical analyses

Data were analyzed using Prism 7.0 software (GraphPad). Clinical EAE data were analyzed by Mann-Whitney *U* test. Other experiments were analyzed with parametric tests (two-tailed Student’s *t* test or one- or two-way ANOVA), with correction for multiple comparisons where appropriate. Colocalization analysis was analyzed with Mander’s Coefficients (M1 and M2) using Fiji software (NIH). A *P* value of less than 0.05 was considered statistically significant. Samples were excluded only if determined to be significant outlier, using Grubb’s test with a significance level of α = 0.01 (GraphPad). Data are expressed as means ± SEM. Sample sizes are indicated in the figure legends.

### Data availability

The data from this study are tabulated in the main paper and supplementary materials. All reagents are available from R.S.K under a material transfer agreement with Washington University. The data that supports the findings of this study are also available from the corresponding author upon request.

### Study approval

All animal experiments followed the guidelines approved by the Animal Care and Use Committee of Washington University School of Medicine. The use of the de-identified post-mortem tissues obtained from a pre-existing repository was not classified as human studies research, per Washington University’s Institutional Review Board guidelines.

## Author contributions

Conceptualization: SM, JLW, and RSK; design of study methodology: SM, JLW and RSK; analysis: SM, JLW, JMB, and LLV; investigation: SM, JLW, LLV, BB, JMB and SA; writing of the original draft: SM and JLW; review and editing of the manuscript: SM, JLW, GFW and RSK; supervision: JLW and RSK.

## Acknowledgements

The authors thank Dr. Angela Archambault for generating T cell lines that were used in adoptive transfer experiments (Washington University in St. Louis, St. Louis, MO). The authors thank Dr. Anne Cross for providing MS tissue specimens and Dr. Robert Schmidt for pathological assessment of MS tissue (Washington University in St. Louis, St. Louis, MO). **Funding**: This study was supported by a postdoctoral fellowship from the National Multiple Sclerosis Society (JLW), National Institutes of Health/National Institute of Allergy and Infectious Diseases grant K22AI125466 (JLW), the National Institutes of Health/National Institute of Neurological Disorders and Stroke Grant F31 NS108629-01A1 (SM), the National Institutes of Health/National Institute of Neurological Disorders and Stroke Grant P01 NS059560 (RSK), R01 NS106289 (GFW), and research grants from the National Multiple Sclerosis Society (RSK and GFW).

## References

1. Reich DS, et al. Multiple Sclerosis. N Engl J Med. 2018;378(2):169–80.

2. Weinshenker BG, et al. The natural history of multiple sclerosis: a geographically based study. I. Clinical course and disability. Brain. 1989;112 ( Pt 1):133–46.

3. Lublin FD, et al. Defining the clinical course of multiple sclerosis: the 2013 revisions. Neurology. 2014;83(3):278–86.

4. Chitnis T, and Weiner HL. CNS inflammation and neurodegeneration. J Clin Invest. 2017;127(10):3577–87.

5. Dendrou CA, et al. Immunopathology of multiple sclerosis. Nat Rev Immunol. 2015;15(9):545–58.

6. Galli E, et al. GM-CSF and CXCR4 define a T helper cell signature in multiple sclerosis. Nat Med. 2019;25(8):1290–300.

7. Wingerchuk DM, and Carter JL. Multiple sclerosis: current and emerging disease-modifying therapies and treatment strategies. Mayo Clin Proc. 2014;89(2):225–40.

8. Baecher-Allan C, et al. Multiple Sclerosis: Mechanisms and Immunotherapy. Neuron. 2018;97(4):742–68.

9. Giles DA, et al. CNS-resident classical DCs play a critical role in CNS autoimmune disease. J Clin Invest. 2018;128(12):5322–34.

10. Keller CW, et al. ATG-dependent phagocytosis in dendritic cells drives myelin-specific CD4(+) T cell pathogenicity during CNS inflammation. Proc Natl Acad Sci U S A. 2017;114(52):E11228–E37.

11. Mundt S, et al. Conventional DCs sample and present myelin antigens in the healthy CNS and allow parenchymal T cell entry to initiate neuroinflammation. Sci Immunol. 2019;4(31).

12. Comabella M, et al. Targeting dendritic cells to treat multiple sclerosis. Nat Rev Neurol. 2010;6(9):499–507.

13. Read SA, et al. Macrophage Coordination of the Interferon Lambda Immune Response. Front Immunol. 2019;10:2674.

14. Yin Z, et al. Type III IFNs are produced by and stimulate human plasmacytoid dendritic cells. J Immunol. 2012;189(6):2735–45.

15. Koltsida O, et al. IL-28A (IFN-lambda2) modulates lung DC function to promote Th1 immune skewing and suppress allergic airway disease. EMBO Mol Med. 2011;3(6):348–61.

16. Dolganiuc A, et al. Type III interferons, IL-28 and IL-29, are increased in chronic HCV infection and induce myeloid dendritic cell-mediated FoxP3+ regulatory T cells. PLoS One. 2012;7(10):e44915.

17. Hemann EA, et al. Interferon-lambda modulates dendritic cells to facilitate T cell immunity during infection with influenza A virus. Nat Immunol. 2019;20(8):1035–45.

18. Lazear HM, et al. Interferon-lambda restricts West Nile virus neuroinvasion by tightening the blood-brain barrier. Sci Transl Med. 2015;7(284):284ra59.

19. Lazear HM, et al. Interferon-lambda: Immune Functions at Barrier Surfaces and Beyond. Immunity. 2015;43(1):15–28.

20. Syedbasha M, and Egli A. Interferon Lambda: Modulating Immunity in Infectious Diseases. Front Immunol. 2017;8:119.

21. Wells AI, and Coyne CB. Type III Interferons in Antiviral Defenses at Barrier Surfaces. Trends Immunol. 2018;39(10):848–58.

22. Sheppard P, et al. IL-28, IL-29 and their class II cytokine receptor IL-28R. Nat Immunol. 2003;4(1):63–8.

23. Sommereyns C, et al. IFN-lambda (IFN-lambda) is expressed in a tissue-dependent fashion and primarily acts on epithelial cells in vivo. PLoS Pathog. 2008;4(3):e1000017.

24. Goel RR, et al. Interferon lambda promotes immune dysregulation and tissue inflammation in TLR7-induced lupus. Proc Natl Acad Sci U S A. 2020;117(10):5409–19.

25. Xu L, et al. Interleukin-29 Enhances Synovial Inflammation and Cartilage Degradation in Osteoarthritis. Mediators Inflamm. 2016;2016:9631510.

26. Won J, et al. Inhaled delivery of Interferon-lambda restricts epithelial-derived Th2 inflammation in allergic asthma. Cytokine. 2019;119:32–6.

27. Dai J, et al. IFN-lambda1 (IL-29) inhibits GATA3 expression and suppresses Th2 responses in human naive and memory T cells. Blood. 2009;113(23):5829–38.

28. Jordan WJ, et al. Human interferon lambda-1 (IFN-lambda1/IL-29) modulates the Th1/Th2 response. Genes Immun. 2007;8(3):254–61.

29. de Groen RA, et al. IFN-lambda-mediated IL-12 production in macrophages induces IFN-gamma production in human NK cells. Eur J Immunol. 2015;45(1):250–9.

30. Greter M, et al. Dendritic cells permit immune invasion of the CNS in an animal model of multiple sclerosis. Nat Med. 2005;11(3):328–34.

31. Simpson JE, et al. Expression of monocyte chemoattractant protein-1 and other beta-chemokines by resident glia and inflammatory cells in multiple sclerosis lesions. J Neuroimmunol. 1998;84(2):238–49.

32. Windhagen A, et al. Expression of costimulatory molecules B7-1 (CD80), B7-2 (CD86), and interleukin 12 cytokine in multiple sclerosis lesions. J Exp Med. 1995;182(6):1985–96.

33. Parker Harp CR, et al. B cell antigen presentation is sufficient to drive neuroinflammation in an animal model of multiple sclerosis. J Immunol. 2015;194(11):5077–84.

34. Nikic I, et al. A reversible form of axon damage in experimental autoimmune encephalomyelitis and multiple sclerosis. Nat Med. 2011;17(4):495–9.

35. Ohsawa K, et al. Involvement of Iba1 in membrane ruffling and phagocytosis of macrophages/microglia. J Cell Sci. 2000;113 ( Pt 17):3073–84.

36. Komiyama Y, et al. IL-17 plays an important role in the development of experimental autoimmune encephalomyelitis. J Immunol. 2006;177(1):566–73.

37. Codarri L, et al. RORgammat drives production of the cytokine GM-CSF in helper T cells, which is essential for the effector phase of autoimmune neuroinflammation. Nat Immunol. 2011;12(6):560–7.

38. El-Behi M, et al. The encephalitogenicity of T(H)17 cells is dependent on IL-1- and IL-23-induced production of the cytokine GM-CSF. Nat Immunol. 2011;12(6):568–75.

39. Rasouli J, et al. Expression of GM-CSF in T Cells Is Increased in Multiple Sclerosis and Suppressed by IFN-beta Therapy. J Immunol. 2015;194(11):5085–93.

40. Caton ML, et al. Notch-RBP-J signaling controls the homeostasis of CD8- dendritic cells in the spleen. J Exp Med. 2007;204(7):1653–64.

41. Clausen BE, et al. Conditional gene targeting in macrophages and granulocytes using LysMcre mice. Transgenic Res. 1999;8(4):265–77.

42. Wang F, et al. Interleukin-29 modulates proinflammatory cytokine production in synovial inflammation of rheumatoid arthritis. Arthritis Res Ther. 2012;14(5):R228.

43. Wu Q, et al. Serum IFN-lambda1 is abnormally elevated in rheumatoid arthritis patients. Autoimmunity. 2013;46(1):40–3.

44. Castillo-Martinez D, et al. Type-III interferons and rheumatoid arthritis: Correlation between interferon lambda 1 (interleukin 29) and antimutated citrullinated vimentin antibody levels. Autoimmunity. 2017;50(2):82–5.

45. Chang QJ, et al. Elevated Serum Levels of Interleukin-29 Are Associated with Disease Activity in Rheumatoid Arthritis Patients with Anti-Cyclic Citrullinated Peptide Antibodies. Tohoku J Exp Med. 2017;241(2):89–95.

46. Srinivas S, et al. Interferon-lambda1 (interleukin-29) preferentially down-regulates interleukin-13 over other T helper type 2 cytokine responses in vitro. Immunology. 2008;125(4):492–502.

47. Morrow MP, et al. Unique Th1/Th2 phenotypes induced during priming and memory phases by use of interleukin-12 (IL-12) or IL-28B vaccine adjuvants in rhesus macaques. Clin Vaccine Immunol. 2010;17(10):1493–9.

48. Egli A, et al. The impact of the interferon-lambda family on the innate and adaptive immune response to viral infections. Emerg Microbes Infect. 2014;3(7):e51.

49. Ye L, et al. Interferon-lambda enhances adaptive mucosal immunity by boosting release of thymic stromal lymphopoietin. Nat Immunol. 2019;20(5):593–601.

50. Hubert M, et al. IFN-III is selectively produced by cDC1 and predicts good clinical outcome in breast cancer. Sci Immunol. 2020;5(46).

51. Blazek K, et al. IFN-lambda resolves inflammation via suppression of neutrophil infiltration and IL- 1beta production. J Exp Med. 2015;212(6):845–53.

52. Gimeno Brias S, et al. Interferon lambda is required for interferon gamma-expressing NK cell responses but does not afford antiviral protection during acute and persistent murine cytomegalovirus infection. PLoS One. 2018;13(5):e0197596.

53. Broggi A, et al. IFN-_λ_ suppresses intestinal inflammation by non-translational regulation of neutrophil function. Nat Immunol. 2017;18(10):1084–93.

54. Espinosa V, et al. Type III interferon is a critical regulator of innate antifungal immunity. Sci Immunol. 2017;2(16).

55. Syedbasha M, et al. Interferon-_λ_ Enhances the Differentiation of Naive B Cells into Plasmablasts via the mTORC1 Pathway. Cell Rep. 2020;33(1):108211.

56. Li Y, et al. Interferon-_λ_ Attenuates Rabies Virus Infection by Inducing Interferon-Stimulated Genes and Alleviating Neurological Inflammation. Viruses. 2020;12(4).

57. Wang T, et al. Toll-like receptor 3 mediates West Nile virus entry into the brain causing lethal encephalitis. Nat Med. 2004;10(12):1366–73.

58. Daniels BP, et al. Viral pathogen-associated molecular patterns regulate blood-brain barrier integrity via competing innate cytokine signals. MBio. 2014;5(5):e01476–14.

59. Ransohoff RM, et al. Three or more routes for leukocyte migration into the central nervous system. Nat Rev Immunol. 2003;3(7):569–81.

60. Li J, et al. Interferon lambda inhibits herpes simplex virus type I infection of human astrocytes and neurons. Glia. 2011;59(1):58–67.

61. Misumi I, and Whitmire JK. IFN-lambda exerts opposing effects on T cell responses depending on the chronicity of the virus infection. J Immunol. 2014;192(8):3596–606.

62. Galani IE, et al. Interferon-lambda Mediates Non-redundant Front-Line Antiviral Protection against Influenza Virus Infection without Compromising Host Fitness. Immunity. 2017;46(5):875–90 e6.

63. Khaitov MR, et al. Respiratory virus induction of alpha-, beta- and lambda-interferons in bronchial epithelial cells and peripheral blood mononuclear cells. Allergy. 2009;64(3):375–86.

64. Vlachiotis S, and Andreakos E. Lambda interferons in immunity and autoimmunity. J Autoimmun. 2019;104:102319.

65. Ye L, et al. Interferon-lambda orchestrates innate and adaptive mucosal immune responses. Nat Rev Immunol. 2019.

66. Prinz M, et al. Distinct and nonredundant in vivo functions of IFNAR on myeloid cells limit autoimmunity in the central nervous system. Immunity. 2008;28(5):675–86.

67. Weber F, et al. Effect of interferon beta on human myelin basic protein-specific T-cell lines: comparison of IFNbeta-1a and IFNbeta-1b. Neurology. 1999;52(5):1069–71.

68. Kozovska ME, et al. Interferon beta induces T-helper 2 immune deviation in MS. Neurology. 1999;53(8):1692–7.

69. Rep MH, et al. Recombinant interferon-beta blocks proliferation but enhances interleukin-10 secretion by activated human T-cells. J Neuroimmunol. 1996;67(2):111–8.

70. Donnelly DJ, et al. Deficient CX3CR1 signaling promotes recovery after mouse spinal cord injury by limiting the recruitment and activation of Ly6Clo/iNOS+ macrophages. J Neurosci. 2011;31(27):9910–22.

71. Kroner A, et al. TNF and increased intracellular iron alter macrophage polarization to a detrimental M1 phenotype in the injured spinal cord. Neuron. 2014;83(5):1098–116.

72. Ank N, et al. An important role for type III interferon (IFN-lambda/IL-28) in TLR-induced antiviral activity. J Immunol. 2008;180(4):2474–85.

73. Baldridge MT, et al. Expression of Ifnlr1 on Intestinal Epithelial Cells Is Critical to the Antiviral Effects of Interferon Lambda against Norovirus and Reovirus. J Virol. 2017;91(7).

74. Cruz-Orengo L, et al. CXCR7 antagonism prevents axonal injury during experimental autoimmune encephalomyelitis as revealed by in vivo axial diffusivity. J Neuroinflammation. 2011;8:170.

75. Lees JR, et al. Regional CNS responses to IFN-gamma determine lesion localization patterns during EAE pathogenesis. J Exp Med. 2008;205(11):2633–42.

76. Williams JL, et al. Targeting CXCR7/ACKR3 as a therapeutic strategy to promote remyelination in the adult central nervous system. J Exp Med. 2014;211(5):791–9.

77. Schindelin J, et al. Fiji: an open-source platform for biological-image analysis. Nat Methods. 2012;9(7):676–82.

78. McCandless EE, et al. CXCL12 limits inflammation by localizing mononuclear infiltrates to the perivascular space during experimental autoimmune encephalomyelitis. J Immunol. 2006;177(11):8053–64.

79. Duong FH, et al. IFN-_λ_ receptor 1 expression is induced in chronic hepatitis C and correlates with the IFN-_λ_3 genotype and with nonresponsiveness to IFN-_α_ therapies. J Exp Med. 2014;211(5):857–68.

