## Supplementary figures and images for "Targeting interferon-_λ_ signaling promotes recovery from central nervous system autoimmunity"

### Supplemental Figure 1

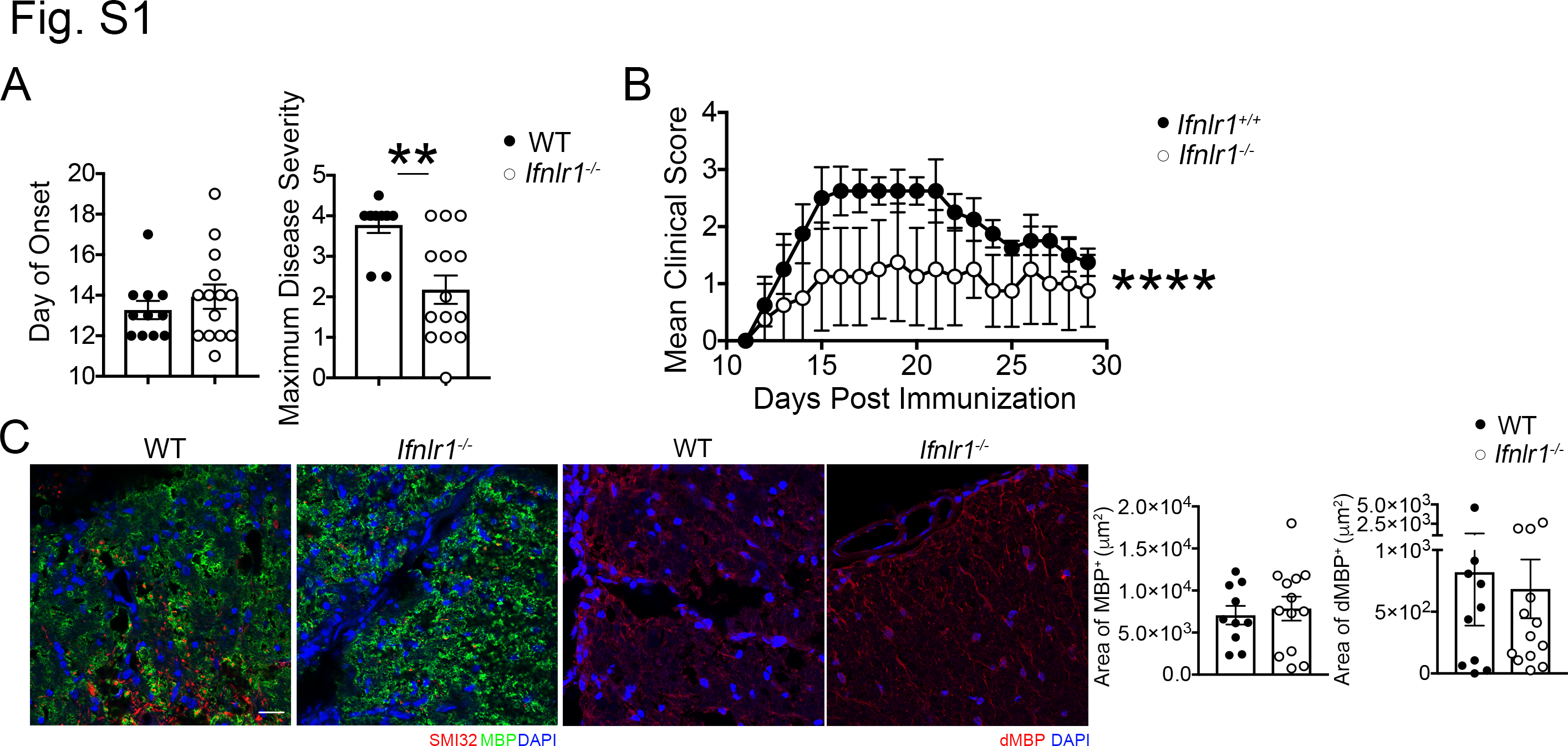

### Supplemental Figure 2

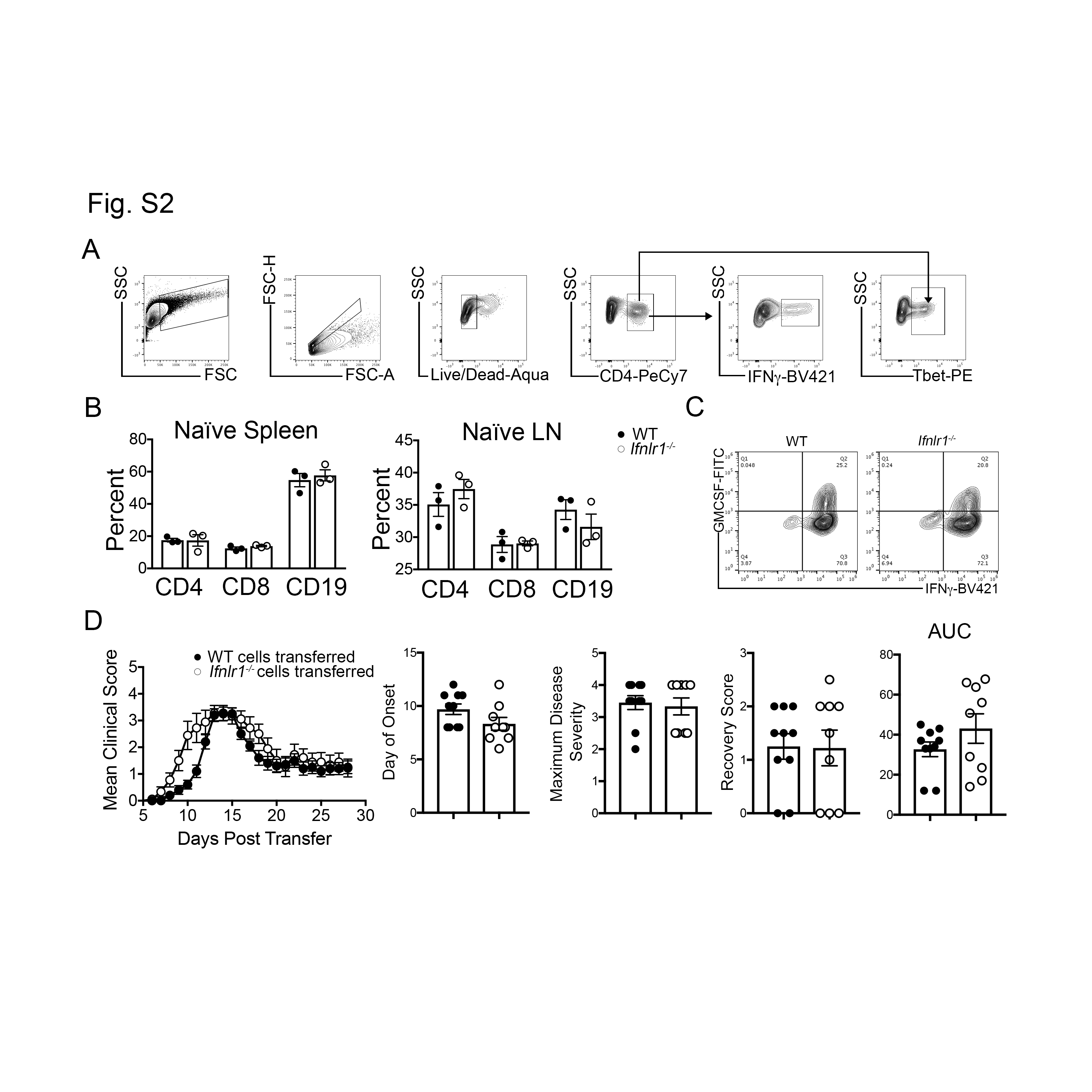

### Supplemental Figure 3

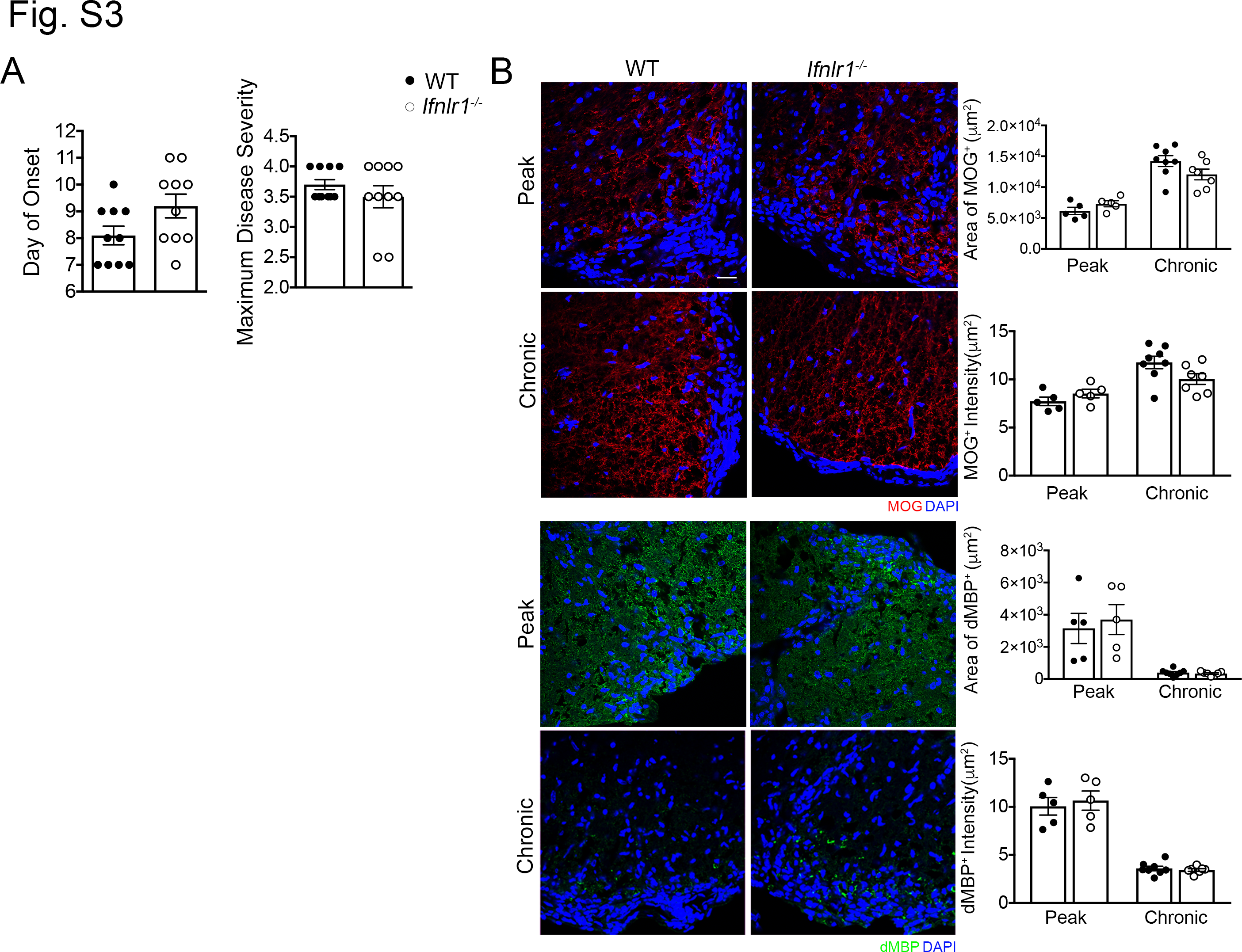

### Supplemental Figure 4

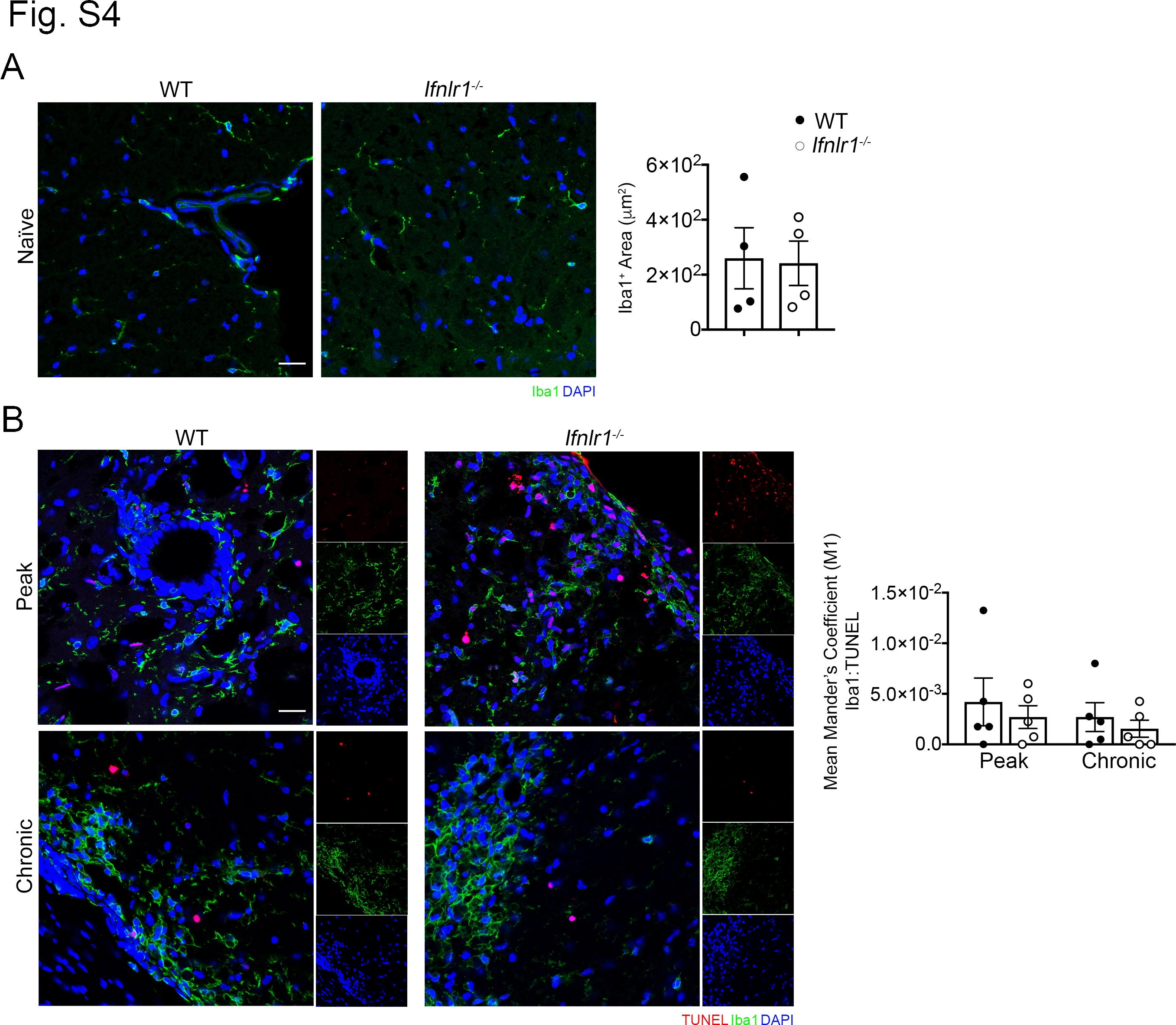

### Supplemental Figure 5

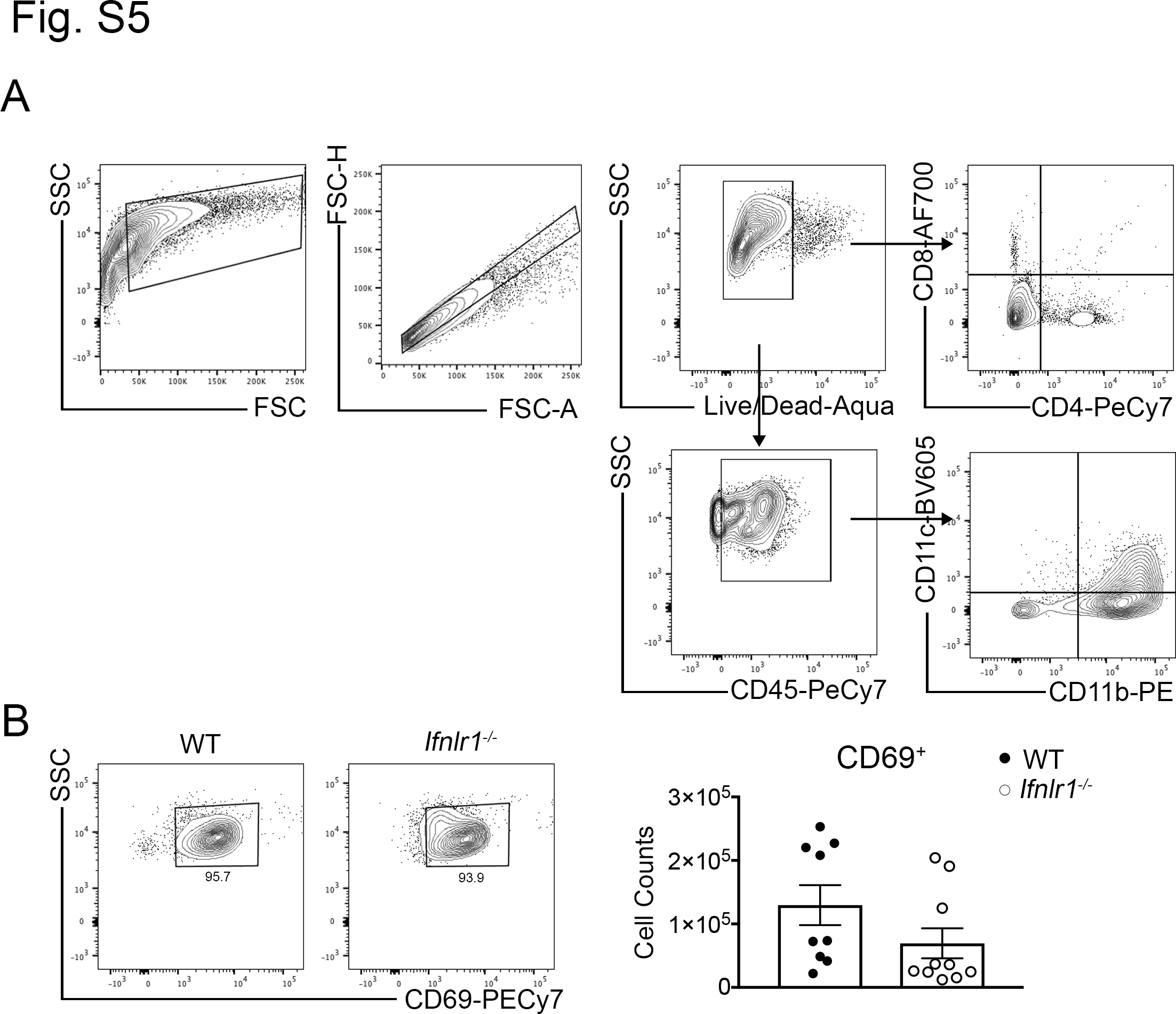

### Supplemental Figure 6

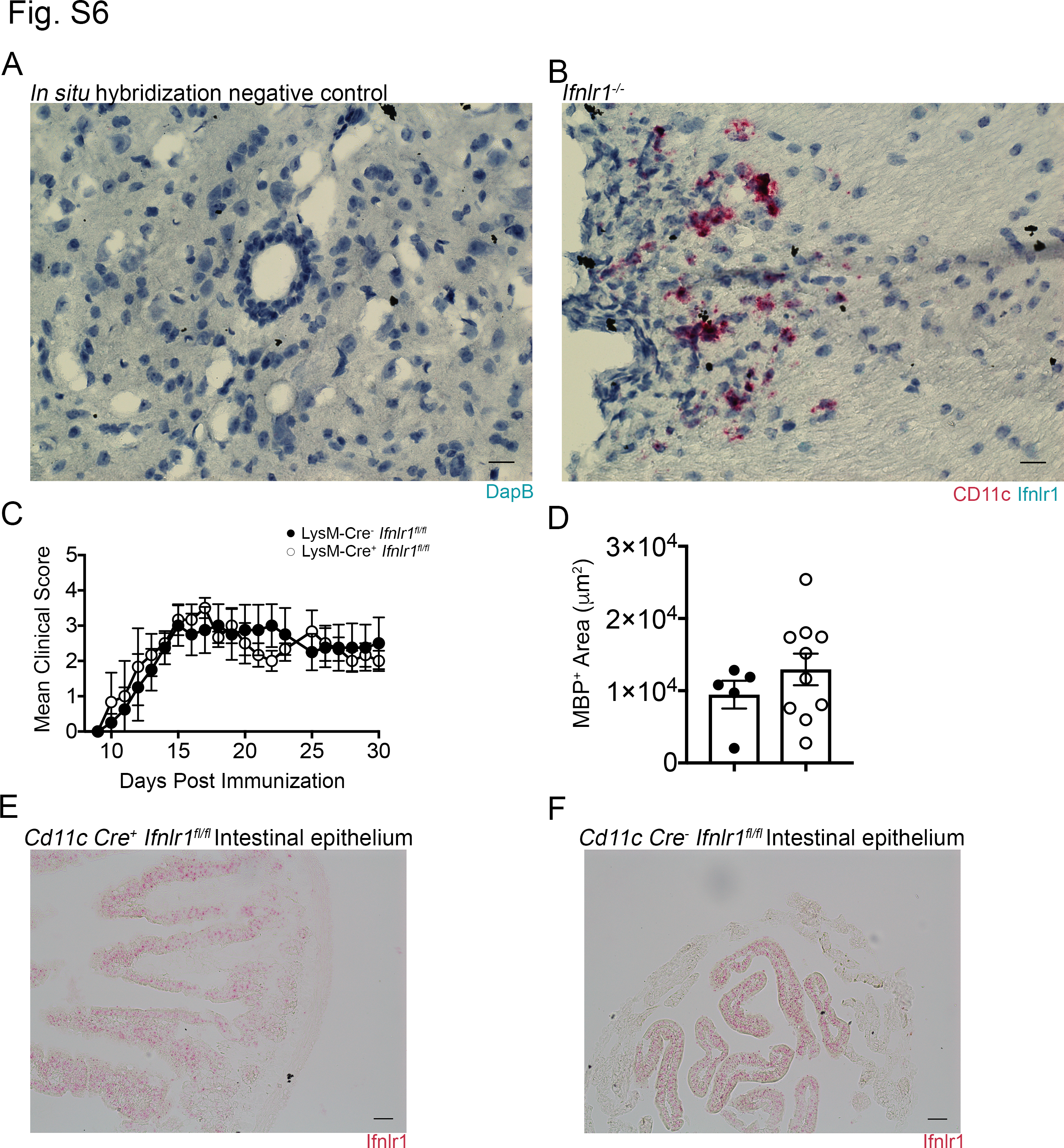
